# Sensory-based quantification of male colour patterns in Trinidadian guppies reveals nonparallel phenotypic evolution across an ecological transition in multivariate trait space

**DOI:** 10.1101/2020.11.23.394668

**Authors:** Lengxob Yong, Darren P. Croft, Jolyon Troscianko, Indar Ramnarine, Alastair Wilson

## Abstract

Parallel evolution, in which independent populations evolve along similar phenotypic trajectories, offers insights into the repeatability of adaptive evolution. Here, we revisit a classic example of parallelism, that of repeated evolution of brighter males in the Trinidadian guppy. In guppies, colonisation of low predation habitats is associated with emergence of ‘more colourful’ phenotypes since predator-induced viability selection for crypsis weakens while sexual selection by female preference for conspicuity remains strong. Our study differs from previous investigations in three respects. First, we adopt a multivariate phenotyping approach to characterise parallelism in multi-trait space. Second, we use ecologically-relevant colour traits defined by the visual systems of the two selective agents (i.e. guppy, predatory cichlid). Third, we estimate population genetic structure to test for adaptive (parallel) evolution against a model of neutral phenotypyc divergence. We find strong phenotypic differentiation that is inconsistent with a neutral model, but only limited support for the predicted pattern of greater conspicuity at low predation. Effects of predation regime on each trait were in the expected direction, but weak, largely non-significant, and explained little among-population variation. In multi-trait space, phenotypic trajectories of lineages colonising low from high predation regimes were not parallel. Our results are consistent with reduced predation risk facilitating adaptive differentiation by female choice, but suggest that this proceeds in (effectively) independent directions of multi-trait space across lineages. Pool-sequencing data also revealed SNPs showing greater differentiation than expected under neutrality and/or associations with known colour genes in other species, presenting opportunities for future genetic study.

## INTRODUCTION

Parallel evolution is defined as the repeated, independent evolution of similar phenotypes under similar selection regimes in multiple independent populations within closely related lineages (Schluter 2004; Bolnick *et al*. 2018). Notable examples include the repeated emergence of stream and lacustrine morphotypes in sticklebacks (McKinnon *et al*. 2004; Schluter *et al*. 2004), and of thin and thick shelled habitat-specific forms of periwinkle (Butlin *et al*., 2014). This phenomenon allows us to ask questions about the predictability of evolution. Does adaptation frequently recapitulate the same phenotypic ‘solutions’ to the same selective challenges? Also, what are the genetic pathways and processes involved? However, characterizing evolutionary trajectories of complex multivariate phenotypes as being either parallel or not is too simplistic. In reality, any two trajectories may be more or less aligned in phenotypic space such that the degree of parallelism lies on quantitative continuum (Stuart *et al*. 2017; Bolnick *et al*.2018). By characterizing the distribution of (multivariate) phenotypes within and among-lineages we can quantify parallelism, testing for similarity in both magnitude and direction of evolutionary trajectories over ecological transitions. We can also assess the importance of parallel (adaptive) evolution within the context of other processes contributing to phenotypic divergence (e.g. drift). Here we apply this approach to a well-known instance of putatively parallel evolution, that of replicated divergence in male colour patterns across predation regimes in the Trinidadian guppy (*Poecilia reticulata*) (Endler *et al*. 1987; Reznick *et al*. 1996).

Animal coloration and pattern traits serve many functions including signalling and crypsis (Cuthill *et al*. 2017) and have long been used to test predictions about the role of predation in driving parallel evolution (Houde, 1997; Allender *et al*. 2003; Steiner *et al*. 2009). High predation risk should select for less ‘conspicuous’ colours and patterns (Endler 1978, 1987; Young *et al*. 2011; Martin *et al*. 2014), but testing this may be sensitive to how colour phenotypes are quantified. Specific colour traits chosen for analysis often vary across studies even within species, while quantitative measures based on human perception (Martin *et al*. 2014) or RGB information (van Belleghem *et al*. 2018; Montenegro *et al*. 2019) may sometimes lack ecological relevance. The latter concern arises because colour signals will (co)evolve with the visual systems (and downstream behaviours) of receiver species (i.e. the sensory drive hypothesis; Endler 1992). For instance, flower traits have coevolved with hymenopteran vision (Dyer *et al*. 2012), and variation in opsin gene sequence and expression is linked to colour polymorphism in African cichlids (Seehausen *et al*. 2008). While this means that human vision could misrepresent colour phenotypes as perceived by relevant selective agents (e.g. conspecific mates, predators), recent advances have improved our ability to model colour variation under different visual models (Stevens *et al*. 2007; Endler *et al*. 2018; Troscianko and Stevens 2015; van den Berg *et al*. 2019). Nevertheless, challenges remain such that colour and pattern phenotypes are necessarily complex and multivariate. Chromatic (e.g. colour) and achromatic (e.g. luminance) aspects of a colour signal are commonly considered, but continued reliance on univariate analyses means that the consequences of trait combinations may be overlooked (Endler and Mappes 2017). Because of these challenges, there has been a call for increased use of spatio-chromatic phenotyping approaches that integrate variation in colour with pattern (spatial arrangement) (Endler and Mielke 2005; Endler *et al*. 2018; van den Berg *et al*. 2019).

A more general limitation of many previous studies has been the relatively infrequent use of population genetic data (but see Steiner *et al*. 2009; Kratochwill *et al*. 2018). Colour traits are important targets of selective processes, but it does not follow that all divergence among populations (whether exhibiting parallelism or not) maps adaptively to local selection regime. Gene flow can sometimes preclude phenotypic divergence between populations despite differences in selection (Räsänen and Hendry 2008; Nosil 2009), while genetic drift could cause (non-adaptive) phenotypic divergence that masks parallelism (Stuart *et al*. 2017; Delisle and Bolnick 2020). Fortunately, patterns of (genome-wide) molecular genetic differentiation can be used to construct null models against which to test for and isolate the phenoptypic signal of local adaptation (e.g., Whitlock and Guillaume 2009; Pascoal *et al*. 2018a,b). There are important caveats to this however, for instance when using phenotypic variation as a proxy for quantitative genetic variation (Pujol *et al*. 2008), or when ‘Isolation by Adaptation’ scenarios are plausible (e.g. Nosil *et al*. 2008; Funk *et al*. 2011). Nonethless, molecular genetic data provide an important opportunity to nuance expectations of phenotypic structuring among populations hypothesised to have undergone parallel evolution. In some cases, notably where dense marker data are available, they can also be used probe the genetic basis of phenotypic differentiation - parallel or otherwise - among populations (Elmer and Meyer 2011; Gautier *et al*. 2018).

In this study, we revisit the well-documented case of putatively parallel evolution of colour in the Trinidadian guppy by combining novel colour phenotyping methods, with estimation of phenotypic divergence among-populations in multivariate trait space and use of population genomic data. Guppies have provided important insights about many evolutionary processes (Endler 1980; Houde 1995; Reznick *et al*. 1997; Magurran 2005) with the highly variable male colour patterns receiving particular attention. Male phenotypes are subject to antagonistic sexual and viability selection; females prefer more ‘conspicuous’ male phenotypes but these also confer greater predation risk. In many rivers in Trinidad, upstream dispersal of piscivores, notably the pike cichlid *Crenacichla frenata*, is limited by barrier waterfalls. Downstream habitats are thus characterised by high predation risk (HP), relative to upstream low predation (LP) sites. Repeated colonization of LP sites has been associated with phenotypic shifts towards brighter, more colourful males (Endler 1983; Millar *et al*. 2006). This is presumably because the costs of being conspicuous (i.e. predation risk) are relatively lower in LP populations while the benefits (i.e. attractiveness to females) remain (Haskins *et al*. 1961; Endler 1980; Houde 1995).

We stress that these patterns of among-population variation in male guppy colour, and their interpretation with respect to predation risk have proven qualitatively robust (Millar *et al*. 2006). However, the diversity of phenotyping methods used across studies has also limited quantitative comparisons of the extent, and direction of (multivariate) evolution across lineages. Phenotyping methods have included scoring the presence/absence, number, size and position of particular colour patches (Endler, 1978; Gotanda *et al*. 2018), spectral measurements (Kemp *et al*. 2019), and spatial pattern approaches (Endler 2012; Endler *et al*. 2018). It is also widely acknowledged that the ecological context is more nuanced than described above (Endler and Houde 1995; Karim *et al*. 2007; Kemp *et al*. 2009). Predation risk varies within HP and LP contexts not just between them (Endler 1995), but canopy cover and light environment also differ between upstream and downstream sites, and frequency-dependent selection also maintains variation within populations (Olendorf *et al*. 2006; Hughes *et al*. 2013). Finally, not all predictions are fully upheld by experiments. For example, low predators risk is thought to allow evolution of larger colourful spots on the body (Haskins *et al*. 1961). However, in a recent study, parallel increases in melanic (black) spots occurred across mesocosms lacking predators, but other colour traits changed inconsistently (Gotanda *et al*. 2018).

Here, we assess the extent of parallel evolution in male guppy colour patterns across repeated instances of LP colonisation from HP ancestors. We implement the Quantitative Colour Pattern Analysis (QCPA) phenotyping approach (van den Berg *et al*. 2019) to model multivariate phenotypic divergence as perceived by the two hypothesized selective agents: namely the guppy (sexual selection) and the Trinidadian pike cichlid (viability selection). Our specific goals are; (i) to determine whether this phenotyping approach recapitulates the expected finding that LP guppies are more ‘conspicuous’ than HP guppies across rivers; (ii) to adopt a geometric perspective (following e.g., Bolnick *et al*. 2018) to quantify the extent of ‘parallelism’ in multivariate trait space; (iii) to evaluate whether the conclusions differ with respect to the two visual systems modelled; and (iv) to assess patterns of phenotypic differentiation (parallel or otherwise) in the context of population genetic structure. We use genomic information obtained via pool-sequencing (hereafter, Pool-seq, Schlötterer *et al*. 2014) to estimate genome-wide population genetic differentiation. The population genomic data also allow us to (v) conduct genome wide scans with genetic regions associated with phenotypic divergence, and thus conduct a preliminary investigation into the genomic basis of colour patterns in the wild.

## MATERIALS and METHODS

### I. Sampling of wild fish

Wild guppies were sampled from 16 sites across 7 rivers in the Northern Mountain Range of Trinidad by seine nets (Table 1) in March 2017. One HP and one or two LP sites were sampled in each river except the Paria (which contributed a single LP site). Fish were transported to the University of West Indies (UWI, St Augustine, Trinidad), housed in 200L aquaria and allowed to acclimate to laboratory conditions for 5-7 days. A subset of males from each population were photographed (Table 1). Subsequently, different individuals (n ∼ 100 per population) from twelve of the 16 sites were shipped live to the University of Exeter (Cornwall, UK). Although primarily for genetic investigations outwith the current study, paternal sibships produced from these wild-caught fish were used in a pilot to validate the present phenotyping strategy (see Supplemental Appendix A for details and results of validation pilot).

**Table 1.** Description of collection site (drainage, river, and regime type) and their GPS coordinates. Sample size denotes fish that were photographed.

| Drainage | River | Regime | Site code | Lat (N) | Long (W) | n<sub>males</sub> |
| --- | --- | --- | --- | --- | --- | --- |
| Caroni | Aripo | high | AH | 10° 39' 02" | 61° 13' 26" | 26 |
| Caroni | Aripo | low | AL | 10° 41' 08" | 61° 13' 56" | 29 |
| Caroni | Aripo | low | AL2 | 10° 41' 10" | 61° 13' 30" | 23 |
| Caroni | Guanapo | high | GH | 10° 39' 28" | 61° 15' 13" | 26 |
| Caroni | Guanapo | low | GL | 10° 42' 42" | 61° 16' 01" | 5 |
| Caroni | Guanapo | low | GL2 | 10° 42' 20" | 61° 15' 48" | 26 |
| Northern | Marianne | high | MH | 10° 45' 54" | 61° 18' 15" | 25 |
| Northern | Marianne | low | ML | 10° 44' 58" | 61° 17' 13" | 23 |
| Northern | Marianne | low | PML | 10° 46' 40" | 61° 18' 04" | 28 |
| Northern | Paria | low | PL | 10° 44' 57" | 61° 15' 54" | 24 |
| Northern | Yarra | high | YH | 10° 47' 23" | 61° 21' 12" | 24 |
| Northern | Yarra | low | YL | 10° 44' 25" | 61° 19' 17" | 26 |
| Oropouche | Quare | high | QH | 10° 39' 53" | 61° 11' 35" | 27 |
| Oropouche | Quare | low | QL | 10° 40' 33" | 61° 11' 49" | 26 |
| Oropouche | Turure | high | TH | 10° 40' 32" | 61° 09' 54" | 26 |
| Oropouche | Turure | low | TL | 10° 41' 31" | 61° 10' 18" | 24 |

### II. Phenotyping of male colour traits

Visible and UV-spectrum photography (Figure 1) were carried out using a Samsung NX1000 camera converted to full spectrum, fitted with a Nikkor EL 80mm lens out on live, unsedated fish housed in a custom designed UV-transparent water filled chamber, as detailed in Appendix A. After photography, fish were humanely euthanized in a lethal dose of buffered MS-222 and tissue preserved in RNAlater (Invitrogen). Photographs were analysed using the QCPA framework, which quantifies colour traits using images and species-specific visual system parameters (see www.empiricalimaging.com and van den Berg *et al*. 2019). In addition to chromatic and achromatic signals, QCPA allows incorporation of spatial information using the geometric arrangement colour patches (with discrimination of adjacent patches based on the receptor noise estimate □*S*; Vorobyev and Osorio 1998). This has ecological relevance because, in behavioral trials, female guppies prefer males that are more conspicuous in colour pattern or have higher discriminant values (Sibeaux *et al*. 2019). We quantified colour variation over the whole body (excluding head, tail and fins), using four complementary traits: saturation boundary strength *sat*_*□S*_, luminance boundary strength *lum*_*□S*_, chromaticity boundary strength *chrom*_*□S*_ and chromaticity boundary strength variation *CoV*_*□S*_ (Endler *et al*. 2018). Together, these measures represent the chromatic (saturation and hue), achromatic (luminance), and spatial (size and position of colour pattern elements) properties of colour patterns. Simplistically, the first three of these can be thought of as describing the “grayness” difference, “brightness” difference, and colour contrast between neighbouring patches respectively. Higher values indicate higher internal contrasts across colour-patch boundaries and/or more boundaries. The final metric describes the variation in colour contrast between patches (i.e. it would be higher if there are both high-contrast *and* low contrast internal boundaries). We used whole body trait measures for two reasons. First, specific pattern elements (i.e. colour patches at particular body locations) differ greatly within and among-populations. Thus, designating universally comparable phenotypes from discrete pattern elements is problematic. Second, female guppies likely select males based on groups of colour patches not single elements (Cole and Endler 2015).

**Figure 1.**
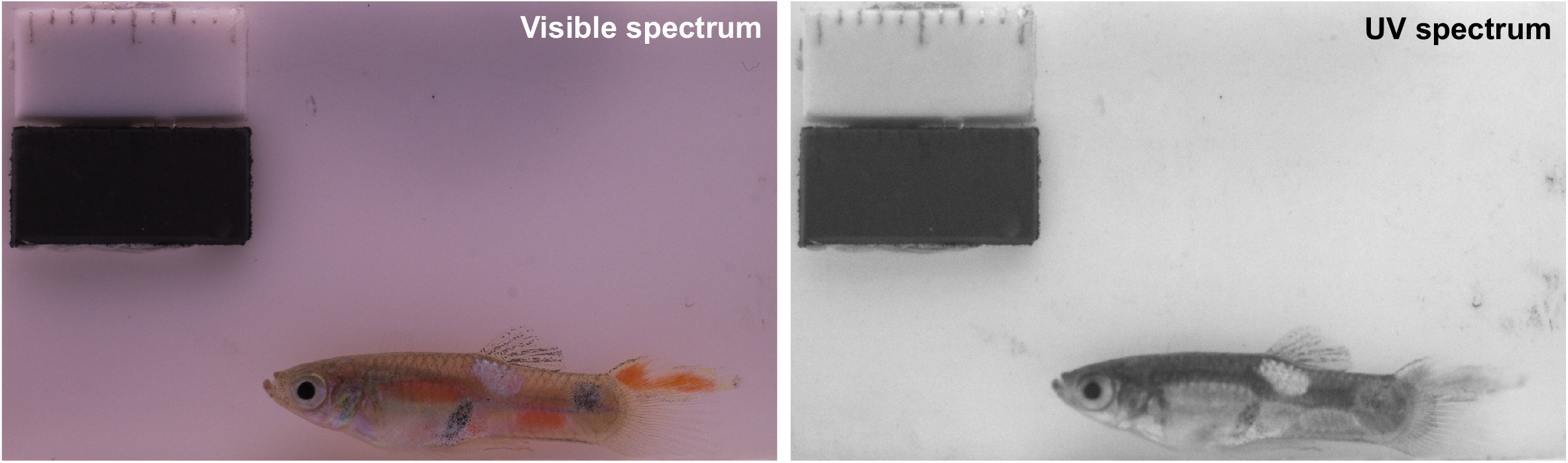
Example of photography setup using calibrated photography using visible and UV spectrum (grayscale). Black and white blocks are 5% and 95% reflectance standards, respectively.

For each photograph, the four traits were scored under two visual system parameter sets; a conspecific (guppy) vision model and a predator (pike cichlid, *Crenacichla frenata*) vision model. Note that *Crenacichla* is used because it is widely viewed as the most important selective visual predator in guppies, although it is not present at the high predation sites sampled in the Yarra and Marianne in this study. The parameters sets included user specified values of (i) receiver spectral sensitivities (the relative abundance of their cones and rods in the retina), (ii) receiver visual acuity, and (iii) physical distance at which the receiver might be from a male guppy. The use of two visual systems reflects our view that perception of male coloration may well differ between receiver species imposing selection (i.e. a pattern that is more conspicuous to a female guppy may not always be so to a cichlid predator).

Support for this possibility was provided by a pilot validation study used to check the suitability of phenotyping methods for detecting genetic variation in male colour and pattern (see Appendix A for full methods and results). Briefly photographs of males from known paternal sibships produced using wild caught parents were processed using our QCPA pipeline. Data analysis confirmed moderate to high sire level repeatabilities (analogous to heritabilities) for all traits under both visual models as expected (since male colour is partly Y-linked; Lindholm and Breden, 2002). This confirms the QCPA approach effectively characterises genetically meaningful phenotypic variation. Importantly, (paternal) family correlations (r_F_) between homologous traits defined under alternative vision models (e.g., *chrom*_*□S*_ based on guppy and cichlid vision models) are positive, but also significantly less than +1 in three of four cases. Using r_F_ as an estimate of the additive genetic correlation (Astles et al. 2006), this indicates homologous metrics defined under the two vision models are genetically distinct (if correlated) traits.

#### III. Generation of Pool-Seq data

To test characterise population genetic structuring we used a Pool-seq approach to obtain allele frequency estimates (Gautier *et al*. 2013; Schlötterer *et al*. 2014). While this approach is cost effective, available funding and tissue samples limited inclusion to 12 of the 16 wild populations (with guppies from the Turure (TH and TL) and Guanapo (GL1 and GL2) rivers excluded). With these exceptions, genomic DNA samples were obtained for 40 fish (20 males; 20 females) per population, using a Qiagen DNAeasy kit (Qiagen Co). For each sample, the concentration and purity of the genomic DNA were measured and using a Nanodrop spectrophotometer and q-bit (Thermo Fisher Scientific). Purity was further checked on a 1% agarose gel before sex-specific DNA pools were created for each population (n = 24; 12 populations by 2 sexes). Pools, each containing equal DNA concentration from 20 individual fish, were sequenced at the Earlham Institute (Norwich, UK). Barcoded DNA paired end libraries with insert size of approximately 150bp were prepared using Illumina Truseq HT library prep and sequenced on 2 lanes using Illumina NovaSeq (Illumina Inc., California, USA). Raw paired-end reads were checked for quality using FastQC, and adapters were verified and removed using *cutadapt*. To investigate among population differences, the raw reads of male and female pools for each population were merged. Reads were then mapped to the reference guppy genome (Guppy_female_1.0+MT, GCA_000633615.2, Künster *et al*. 2016) with BWA mem (Li and Durbin 2010) using default parameters to generate initial BAM files. Aligned reads were then sorted, marked for duplicates and indexed, to generate final BAM files using Samtools v1.9 (Li *et al*. 2009). All BAM files were merged to create a mpileup file (i.e. samtools mpileup –f), which was subsequently filtered for indels and then used to generate a sync file (mpileup2sync.jar; base quality > 20 or --min-qual 20) containing allelic frequency information for every population using *Popoolation2* (v1.201) (Kofler *et al*. 2011). A clean sync file was then obtained after indel removal (filter-sync-by-gtf.pl), and subsequently used for downstream population genomic analyses.

### IV. Data analysis

The phenotypic data set comprised measures of 8 response variables (*sat*_*□S*_, *lum*_*□S*_, *chrom*_*□S*_, *CoV*_*□S*_ x 2 visual system models) for 388 male guppies representing 16 distinct populations (6 HP and 10 LP) from 7 rivers. The genomic data comprised of 24 pools of DNA-seq data (12 populations x 2 sexes), whose summary statistics are described in the Supplemental (Table S1). In brief, each pool generated between 362-599M reads with a mean Phred score of 35.8 and 96x depth of coverage. Between 88-98% of raw reads were successfully mapped to the guppy female reference genome assembly. Unless otherwise stated, all statistical analysis was done in R (R Core Team 2019) with ASReml-R (Butler *et al*. 2017) used to fit mixed models. Full R codes and citations to all packages, bash scripts, and Javascripts for colour analysis will be provided in the supplemental materials upon publication.

#### Testing effects of population and predation regime

We first plotted trait means (by river and overall), to check whether differences between populations and or predation regimes (HP versus LP) were visually apparent. We then fitted univariate mixed models to each trait, including a random effect of *population* (i.e. sampling site) and a fixed effect of predation *regime* (as a two level factor). This allowed us to estimate variance among *populations* (V_Pop_) conditional on the fixed effect of *regime* (i.e. as the among-population variance not explained by *regime*). We scaled traits to standard deviation units before analysis such that V_Pop_ can be interpreted as an intra-class correlation unadjusted for fixed effects. We also estimated the adjusted population level repeatability (R_Pop_) for each trait (i.e., V_Pop_ as a proportion of total variance conditional on the fixed effect). Statistical inference on the fixed effect was by Wald F tests and we used likelihood ratio tests (LRT) to assess the significance of the random effect. For the latter, we assumed that twice the difference in log-likelihoods between full and reduced (with no random population effect) is distributed as a 50:50 mix of χ^2^_0_ and χ^2^_1_ (subsequently denoted χ^2^_0,1_; Stram and Lee 1994).

#### Multivariate models and estimation of pairwise phenotypic distance

We built a multivariate mixed model of all 8 traits. A random effect of population was included on each trait but with no fixed effects (except intercepts). This yielded an estimate of the among-population variance-covariance matrix (**D**), containing among-population trait variances (but no longer conditional on any *regime* effects) on the diagonal, as well as covariances that were scaled to among-population correlations (r_Pop_) between all trait pairs. We assume approximate 95% CI of r_Pop_±1.96 SE.

We then used the fitted multivariate model to predict pairwise phenotypic distances between multivariate population means by extracting the best linear unbiased predictions (BLUP) of trait means by population and calculated the 16×16 (since n_population_=16) Euclidean distance matrix (**E**) among populations in n_trait_-dimensional trait space. We used all 8 traits (**E**_all_), but also generated the matrices based on guppy (**E**_gvm_) and cichlid (**E**_cvm_) vision model traits respectively. Note that uncertainty in BLUP is not reflected in the point estimates of **E**_all_, **E**_gvm_ and **E**_cvm_, but we use these matrices to visualise patterns of among-population divergence not as a basis for robust statistical inference. Specifically, neighbour-joining trees based on **E**_all_, **E**_gvm_ and **E**_cvm_ were plotted to assess any tendency to cluster by *regime*. For each population pair, we also plotted the distance based on the guppy vision model against the corresponding cichlid distance to assess whether (i) a relationship between **E**_gvm_ and **E**_cvm_ is apparent, and ii) phenotypic distances tend to be lower for within-versus across-*regime* population pairs.

#### Testing for association of phenotypic and genetic distance

For the subset of 12 populations with molecular as well as phenotypic data, we used the sync file to estimate genome-wide differentiation among populations using the fixation index (F_ST_). F_ST_ was calculated for single nucleotide polymorphisms (SNPs) in 50k windows using *fst*.*sliding*.*pl* implemented in *Popoolation2* (parameters: –min-count 2 –min-coverage 4 – max-coverage 80 –pool-size 80 –window-size 50000 –step-size 50000), which calculates F_ST_ following Hartl and Clark (1997). The resultant among-population F_ST_ matrix was compared to phenotypic distance estimates in two ways. First, we used Mantel tests to assess the correlation with phenotypic distance (represented by **E**_all_, **E**_gvm_ and **E**_cvm_ sub- setted to the 12 populations with SNP data). Second, following Pascoal *et al*. (2017) we refitted univariate mixed models to each of the 8 response variables, but this time with no fixed effects and a modified random effect structure to total V_Pop_ (i.e. not conditional on *regime*) into two components: one attributable to divergence under neutral processes (i.e. drift; V_Pop.N_); and another attributable, under some assumptions, to divergence under selection (from predation, sexual selection and/or unknown selective agents; V_Pop.S_).

This second approach, follows the same premise as Q_ST_-F_ST_ comparisons (Leinonen *et al*. 2013), and asks whether there is more quantitative phenotypic divergence among-populations than expected from levels of (putatively) neutral molecular divergence. To implement it, we ran models with two random effects of population identity, one which covaries among populations according to their molecular ‘relatedness’ structure, and a second which is uncorrelated across populations. For the former we assume phenotypic covariance between fish sampled from populations *i* and *j* that is attributable to neutral population effects is equal to S_ij_·V_Pop.N_ where S_ij_ is genome-wide ‘similarity’ between the populations. Since F_ST_ inscreases wih dissimilarity we define 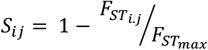 where 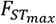 is the highest observed pairwise F_ST_ among the populations sampled. This simply scales *S*_*ij*_ from a maximum of 1 (when i=j) to 0 (for the most differentiated pair of populations). To test whether trait variation among populations is greater than expected under neutral divergence alone, we compared the model with both variances to one where V_Pop.S_ is absent using LRT. Two caveats to this method shoud be noted: first, it assumes that to a first approximation genome-wide F_ST_ measures neutral differentiation (i.e. from drift and gene flow); second, since we are modelling phenotypic variation in wild caught fish rather than in a common-garden experiment, any contributions to population divergence from phenotypic plasticity are likely to be partitioned into V_Pop.S_ (Pujol *et al*. 2008).

#### Multivariate models for estimating parallelism among replicate population pairs

Phenotypic analyses described above are ‘blind’ to the ancestral versus derived status of HP and LP population pairs with rivers. Therefore, we also adopted recent methods that use phenotypic change within lineages to quantify parallelism (De Lisle and Bolnick 2020). We assume that within each river, an HP and LP pair can be viewed as representing a ‘lineage’ in which the HP is ancestral. This assumption yielded 9 ‘lineages’ among the 16 populations with phenotypic data. Fish from the Paria are excluded as there is no HP site, but the Aripo, Guanapo, and Marianne rivers each contribute two ‘lineages’ (i.e. HP-LP comparisons). We describe the approach briefly, using notion from De Lisle and Bolnick (2020) and referencing their equation numbers. First we calculated, the n_trait_ row by m_lineage_ column data matrix **X**, where each element of **X** represents Δ*z*_n,m_ (the difference in mean standardized phenotype (z) between HP and LP populations for trait n in lineage m; Equation 3). Each row of **X** thus represents the vector of phenotypic change from HP to LP within a lineage in multi-trait space. **X** was used to calculate **C**, the (m x m) among-lineage correlation matrix of phenotypic change vectors which was then subject to eigen decomposition (following Equations 5 and 6). To mirror our investigation of the among-population distance matrices (i.e. **E**_all_, **E**_gvm_ and **E**_cvm_), we calculated and decomposed **C** matrices based on *sat*_*□S*_, *lum*_*□S*_, *chrom*_*□S*_ and *CoV*_*□S*_ determined using both vision models (**C**_all_), but also using guppy (**C**_gvm_) and cichlid vision model (**C**_cvm_) traits, separately.

In the case that multivariate trajectories are perfectly parallel across all lineages, the first eigen vector of **C** should explain all variation and all lineages would load on this vector with the same sign (but differing magnitudes if the extent, rather than direction, of phenotypic change differed among lineages). In contrast, a uniform distribution of eigen values (meaning low among-lineage correlations of change vectors), and/or a mixture of positive and negative lineage-specific loadings on the first eigen vector is expected under non-parallelism. As suggested by De Lisle and Bolnick (2020), for the case of fewer traits (here 4 or 8 depending on the version of **C**) than lineages (here 9), we generated a null distribution of *m* eigenvalues by simulating 1000 random vectors representing evolutionary change under independence (among lineages) of multivariate phenotypic trajectories. While acknowledging that power to reject the null (no parallelism) may be low (De Lisle and Bolnick 2020), observed eigenvalues were then compared to the simulated distribution. An eigenvalue of **C** greater than 95% of values simulated assuming independence is taken as statistical support for parallelism.

#### Genotype-phenotype association mapping from pool-seq data

Leveraging the pool-seq data, we tested for relationships between allele frequencies at SNP loci and phenotypic variation (colour metrics: *sat*_*□S*_, *lum*_*□S*_, *chrom*_*□S*_ and *CoV*_*□S*_) and predation regime (*high* vs *low*). We did this using genome-wide scans for association in BayPass (Gautier, 2015). This method identifies loci more with higher than expected differentiation among populations based on the *XtX* statistic (Gunther and Coop 2013), and alsotests associations between SNPs and population-specific covariables (while accounting for background population structure from drift and gene flow). Since direct selection on male coloration is necessarily sex-limited, we elected to use SNP data from male pools only for these analyses (rather than combined male and female data used for genome wide F_ST_ estimates). Major and minor allele counts for each SNP were calculated for each population from the previously compiled sync file using the *snp-frequency-diff*.*pl* script with the following parameters (–min-coverage 75 –max-coverage 200). Using custom awk scripts, allele counts were extracted and formatted for BayPass’s input genotype file (Gautier 2015). The large number of calculated SNPs from the male pools (ca. 5,977,803 SNPs) mainly on known linkage groups (LGs) were then subsampled to yield about 100k SNPs along the whole genome (Mean = 95,911 ± 61.2 SNPs from known LGs only, corresponding to about 4170 per LG) to generate about 100 sub-datasets. Subsets were then analysed individually using the BayPass core and standard covariate models. The resulting population covariance matrices of allele frequencies (interpretable as similarity matrices) were generated under the core model and checked for consistency across all data subsets by calculating the distance among matrices (hereafter FMD) using the *fmd*.*dist* function (Figure S1; Table S2). The mean FMD among all pair wise comparison was 0.2 (with r > 0.99 for all pairwise comparisons).

To identify SNPs that deviated from neutral expectations we used *simulate*.*baypass*. This uses the population covariance matrix to simulated ‘pseudo-observed datasets’ (POD), assuming SNP neutrality, consistent with the demographic history. This allows a null distribution of the *XtX* to be generated, the 99% quantile of which was used as a significance threshold (median *XtX* > 21.2) to determine whether observed SNPs were under selection. We then used the BayPass IS covariate models, and resultant Bayes Factor (BF) to test for association between SNPs and population level male colour traits (and/or predation regime) (Gunther and Coop 2013; Gautier 2015). Evidence for significant association between SNP and a tested covariate was based on BF > 30db (Gautier 2015).

Finally, we checked whether SNPs identified as deviating from neutrality and/or associated with a covariate showed enrichment for gene ontology (GO) categories and functions. We extracted the corresponding identity of genes or nearby genes, which and blasting against the zebrafish genome (*Danio rerio* NCBI Refseq GCF_000002035.6, Howe *et al*. 2013) for annotated orthologs. The annotated orthologs were run in the gene ontology resource (http://geneontology.org/, Ashburner *et al*. 2000; The Gene Ontology Consortium 2019).

## RESULTS

### I. Non-significant predation regime effect on and among-population variance in male colour patterns

Mean values of *chrom*_*ΔS*_, *cov*_*ΔS*_, *sat*_*ΔS*_ and *lum*_*ΔS*_ varied considerable between vision models and populations (Figure 2a-d). Qualitatively, the guppy vision model detected less variability among populations and tended to yield higher (more conspicuous) values for *chrom*_ΔS_ and sat_ΔS_ relative to cichlid vision (with the converse for *cov*_*ΔS*_ and *lum*_*ΔS*_). Differences between HP and LP regime were not consistent across traits and rivers, although overall (i.e. across rivers) trait means were higher in LP guppies for all 8 traits.

**Figure 2.**
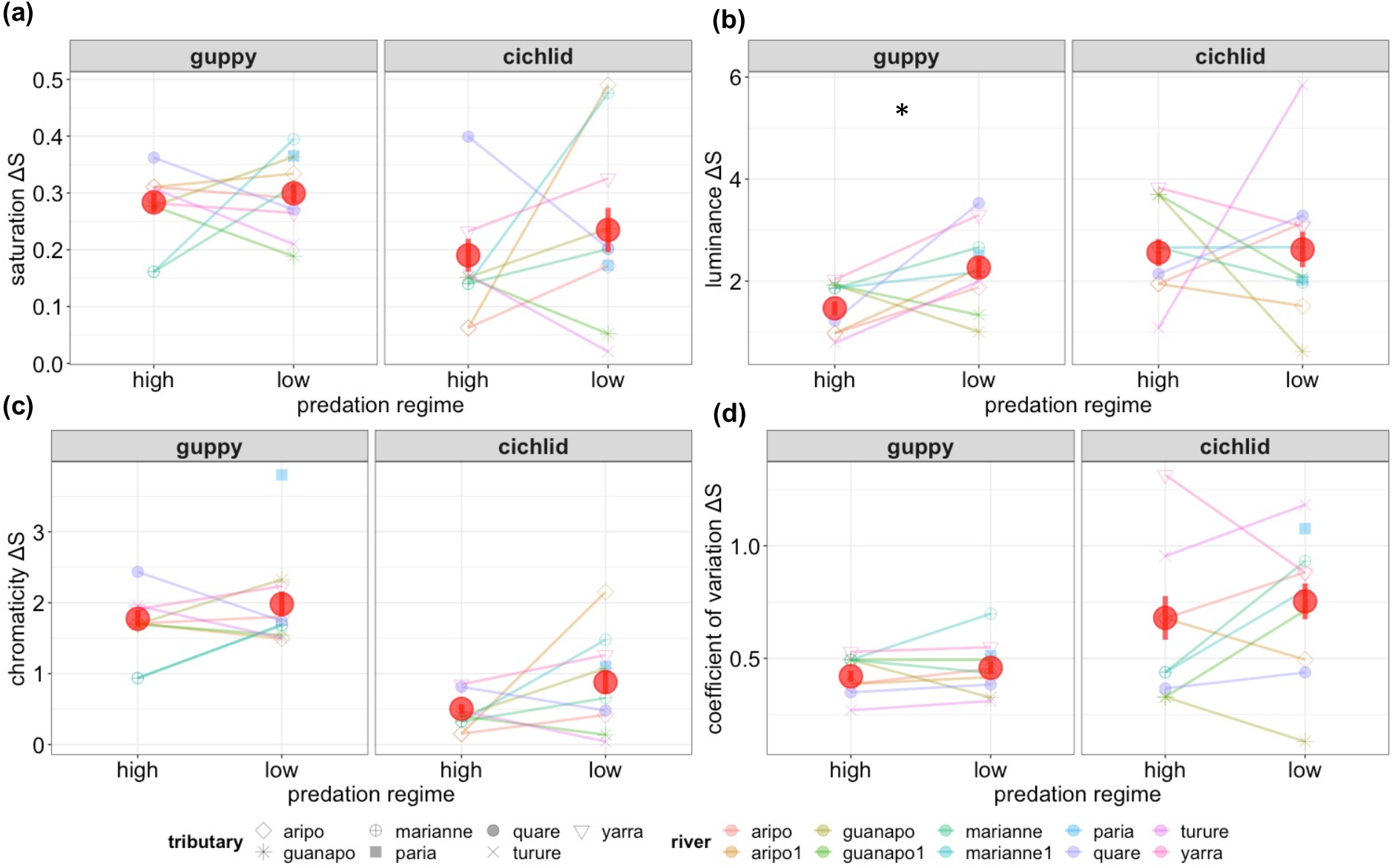
Distribution of population mean trait values (prior to conversion to SDU) by predation regime and population under guppy and pike cichlid vision models for (a) colour saturation (*sat*_*ΔS*_), (b) luminance (*lum*_*ΔS*_), (c) chromaticity (*chrom*_ΔS_), (d) coefficient of variation in chromaticity (*chrom*_*ΔS*_). Lines link populations of differing regime within rivers. Also shown are the overall trait means by predation regime (red circle) with standard error. Asterisk (*) denotes significant difference in overall mean between regimes. Note that Paria (light blue square) does not have a corresponding high predation population.

Univariate mixed models confirmed this pattern, but provided limited statistical support for differences in trait means between HP and LP (Table 2). Positive *regime* coeffcients for all traits indicated a tendency to higher conspicuousness at LP sites; but across the eight models, only *lum*_*ΔS*_ under the guppy vision model was significant (coefficient (SE): 0.83 (0.37) sdu, P < 0.03; Table 2). A moderately large yet non-significant effect (+0.57 sdu) was estimated for chrom_ΔS_ under the cichlid model, while the average estimated *regime* effect size of *regime* +0.328 sdu. Support for strong among-population differentiation, over and above any *regime* effects was unequivocal; LRT comparisons to reduced models with no population effect yielding P<0.001 for all traits (Table 2). Population level (conditional) repeatability R_Pop_ ranged from 30% to 71% with an average of 54%. R_Pop_ estimates were very close to V_Pop_ indicating that *regime* explained little of the among-population differentiation (if *regime* explained large amounts of trait variance, then in these models V_Pop_ + V_R_ would <1 leading to a systematic finding of R_Pop_>V_Pop_).

**Table 2.**
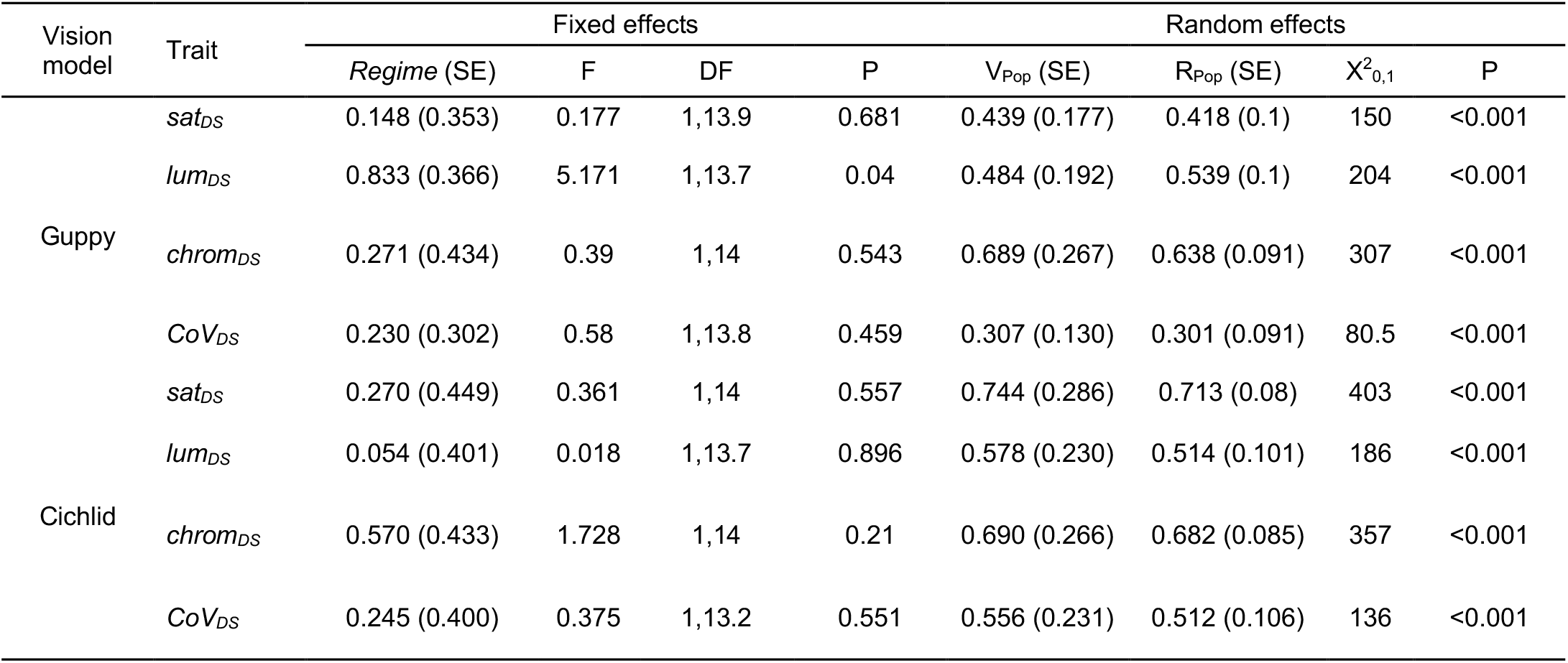
Estimated regime (fixed) and population (random) effects from univariate mixed models of each trait. Regime effects denote the consequence of low relative to high predation regime with inference by F-tests. Among-population variance (V_Pop_) is conditional on fixed effects and was tested by likelihood ratio test. Also shown is the population level repeatability (R_Pop_).

### II. No clustering of guppy colour patterns by predation regime

The **D** matrix (estimated without conditioning on *regime*) yielded V_Pop_ estimates very similar to (conditional) estimates obtained from univariate models (Table 3). The correlation structure within **D** was not universally positive (which is the expectation if populations varied along a simple axis from greater to less conspicuousness; Table S3). Population-level correlations between homologous traits defined using guppy and cichlid vision models were positive but significantly less than +1 (assuming an upper 95% confidence interval of r_Pop_ + 1.96 SE). Specifically r_pop_ (SE) were estimated as 0.65 (0.16), 0.39 (0.23), 0.28 (0.24) and 0.09 (0.28) across the two models for *sat*_*ΔS*_, *lum*_*ΔS*_, *chrom*_*ΔS*_, and *CoV*_*ΔS*_ respectively.

**Table 3.** Estimates of among-population repeatabilities (R_Pop_) derived from univariate mixed models that partition total population effects into components attributable to putative neutral (R_Pop.N_) versus selective (R_Pop.S_) processes. Also shown are LRT comparisons to a reduced model in which all among-population variance is assumed to have a neutral basis. Note where a variance component, and so the corresponding repeatability, was bound to zero to keep it in allowable parameter space no estimate of the SE is possible.

| Vision model | Trait | $R_{Pop}$ (SE) | $R_{Pop,N}$ (SE) | $R_{Pop,S}$ (SE) | $X^2_{0,1}$ | P |
| --- | --- | --- | --- | --- | --- | --- |
| Guppy | *sat<sub>DS</sub>* | 0.385 (0.109) | 0.000 (-) | 0.385 (0.109) | 3.787 | 0.026 |
|  | *lum<sub>DS</sub>* | 0.414 (0.144) | 0.073 (0.083) | 0.342 (0.196) | 4.537 | 0.017 |
|  | *chrom<sub>DS</sub>* | 0.654 (0.136) | 0.031 (0.092) | 0.623 (0.208) | 4.95 | 0.013 |
|  | *cov<sub>DS</sub>* | 0.240 (0.089) | 0.000 (-) | 0.240 (0.089) | 9.651 | 0.001 |
| Cichlid | *sat<sub>DS</sub>* | 0.418 (0.109) | 0.418 (0.109) | 0.000 (-) | 0 | 0.5 |
|  | *lum<sub>DS</sub>* | 0.364 (0.108) | 0.000 (-) | 0.364 (0.108) | 5.286 | 0.011 |
|  | *chrom<sub>DS</sub>* | 0.400 (0.108) | 0.400 (0.108) | 0.000 (-) | 0 | 0.5 |
|  | *cov<sub>DS</sub>* | 0.495 (0.114) | 0.000 (-) | 0.495 (0.114) | 8.581 | 0.002 |

Phenotypic distance matrices derived from the multivariate mixed model fit provided little support for clustering of populations by *regime* in multi-trait space (Figure 3, Table S4). Using distance defined from all traits at once and guppy vision models, there was arguably some patterning (e.g., upper right portion of Fig 3a contains a grouping of 6 LP populations with 1 HP whereas Fig 3b has a grouping of 6 LP and no HP), but this is less apparent using just the traits subsets from the predator or cichlid (Figure 3c) vision model.

**Figure 3.**
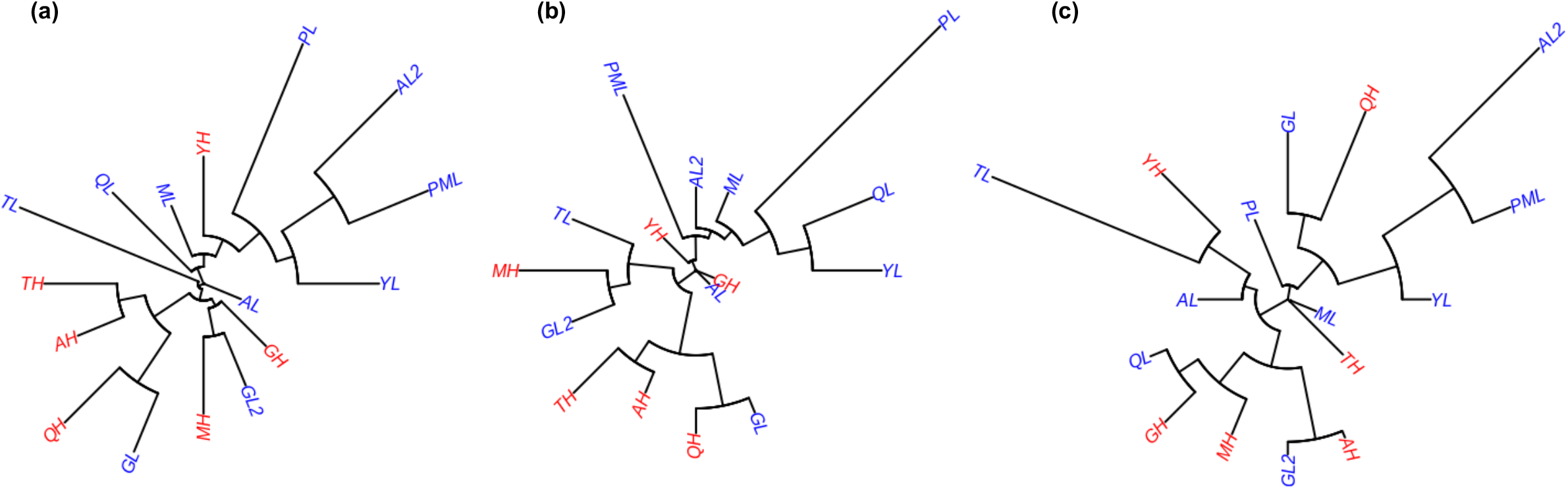
Neighbour joining trees of populations based on estimated phenotypic distance matrices in n-dimensional trait space. Tree shown are determined using (a) all eight colour measures, (b) the four traits derived from the guppy-and (c) the four traits derived from the cichlid-vision model. Red and blue labels denote high and low predation *regime* populations, respectively.

Pairwise phenotypic distances (using either guppy vision or cichlid vision traits) did not suggest shorter average pairwise distances between populations within-versus across-predation regimes (Figure 4). If clustering by *regime* was present, we would expect higher phenotypic distances for low-high predation (LH) comparisons than for low-low predation (LL) or high-high predation (HH) (i.e. the ellipse of red LH points in Figure 4 would be shifted to higher values on x and/or y axes than the blue ones; Figure 4). Two further points emerge from Figure 4; first, there is no apparent relationship between the pairwise population distance estimates defined by the two vision models. Population pairs that are more distinct with respect to guppy vision are not generally more distinct under the cichlid vision model. This is consistent with the absence of strong positive r_POP_ between homologous traits noted above. Second, the 95% confidence ellipse of pairwise distances for HH comparisons is smaller and shifted to lower values (on both axes) relative to HL or LL comparison. This pattern implies that there may be lower phenotypic variance among high predation populations that among low predation populations. To investigate further, we fitted post hoc multivariate mixed models (described Table S5) to regime specific data subsets, and compared total phenotypic variance among-populations (calculated as the trace of regime specific **D** matrices). Point estimates corroborated our interpretation of Figure 4, the trace of **D**_**H**_ being 65% of the trace of **D**_**L**_. However, unsurprisingly given the number of populations in each regime, our trace estimates were characterized by high uncertainty and the difference is not statistically significant.

**Figure 4.**
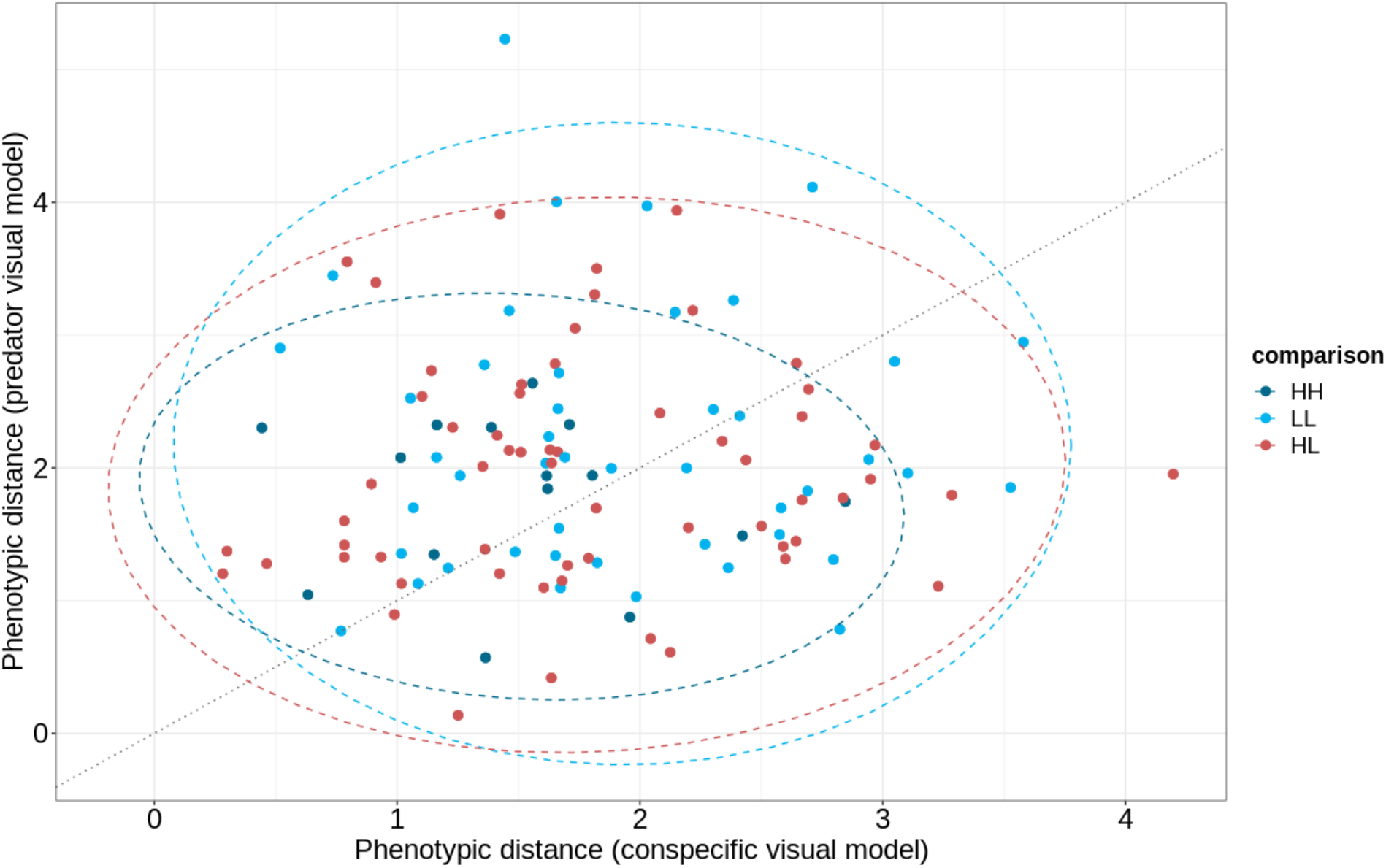
Between-population distance estimates based on guppy (x-axis) and cichlid (y-axis) vision models. Each point denotes a single pairwise population comparison. Red points denote cross-regime distances between a high and a low predation site and blue points denote pairwise distances within predation regimes (dark blue for high, light blue for low), Ellipses illustrate approximate 95% confidence limits of the distributions and the 1:1 line is also shown.

### III. Phenotypic differentiation does not align with putative neutral genetic divergence

Consistent with previous studies (e.g., Suk and Neff 2009; Willing *et al*. 2010; Fraser *et al*. 2015), genome-wide pairwise F_ST_ comparisons revealed strong genetic structuring by river and drainage (Figure 5; Table S6). There was almost no correlation between molecular differentiation and estimated phenotypic distance (Mantel test of F_ST_ matrix correlation with **E**_all_ r=-0.049, P=0.608; with **E**_gvm_ r=0.026, P=0.426; and with **E**_cvm_ r=-0.033, P=0.569). A neutral model of among population divergence was also rejected (at α=0.05) for 6 out of 8 traits using our mixed model strategy to partition V_Pop_ into components attributable to neutral (V_Pop.N_) and selective (V_Pop.S_) components (Table 3). The proportion of trait variance explained by the latter was considerably higher than by the former for most traits. Under the cichlid vision model, *sat*_*ΔS*_ and *chrom*_*ΔS*_ proved exceptions to this pattern, with the more complex model offering no improvement to model fit and V_Pop.S_ being bound to zero. Thus, with these two exceptions, genome wide-molecular divergence among populations cannot readily explain the phenotypic differentiation structure.

**Figure 5.**
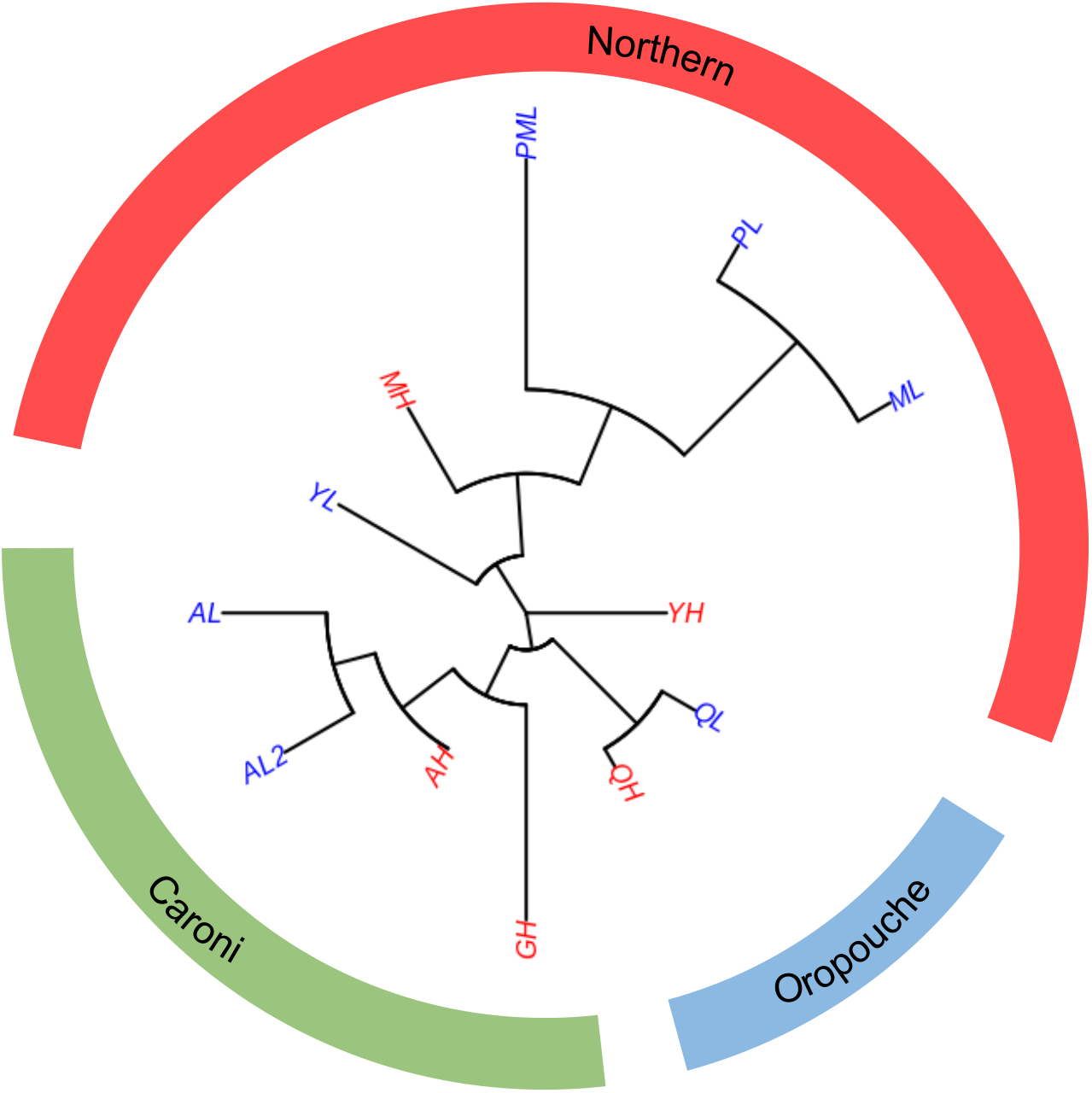
Neighbor joining tree based on whole-genome population F_ST_ distance, clustering by rivers, then drainages, consistent with the independent evolution of each replicate. Branch length represents Fst. Populations: AH = Aripo High; AL = Aripo Low; AL2 = Aripo Low 2; GH = Guanapo High; MH = Marianne High; ML = Marianne Low; PML = Petite Marianne; PL = Paria; QH = Quare High; QL = Quare Low; YH = Yarra High; YL = Yarra Low. Blue denotes Low predation regime, whereas Red High.

### IV. Non-parallelism in guppy colour pattern among lineages

Decompositions of the three **C** matrices revealed a highly skewed distribution of eigenvalues (Table 4). However, since the rank of **C** is limited to the smaller of n (traits, here either 4 or 8) and m (lineages, here 9) all three 9 × 9 **C** matrices will necessarily be rank deficient. In this context, eigen values of zero are inevitable and not biologically informative. Moreover, the first eigenvectors were not particularly dominant, capturing 55%, 51% and 40% of **C**_gvm,_ **C**_cvm_ and **C**_all_ respectively, while lineages loaded on this first eigenvector with a mixture of positive and negative signs. The structure of the **C** matrices was thus not consistent with parallel phenotypic evolution among these lineages in 8-trait space, or in either of the 4-trait spaces. Random vectors provided no evidence that the first eigen values were larger than expected under a null model of independent evolutionary trajectories across lineages (Figure S2). The second eigen values of **C**_gvm_ and **C**_all_ were greater than 95% of simulated values, which suggested some overdispersion and thus deviation from complete independence. However, lineages also loaded on this second eigenvector with a mix of signs too (in all three **C** matrices) so we cannot interpret this as parallel evolution in a direction defined by the second axis of **C**. We suspect that this signal of non-independence arises because data from single high predation populations in Aripo, Guanapo, and Marianne rivers each contributed to the vectors of phenotypic change for two lineages (i.e. HP vs LP comparisons within-river).

**Table 4.**
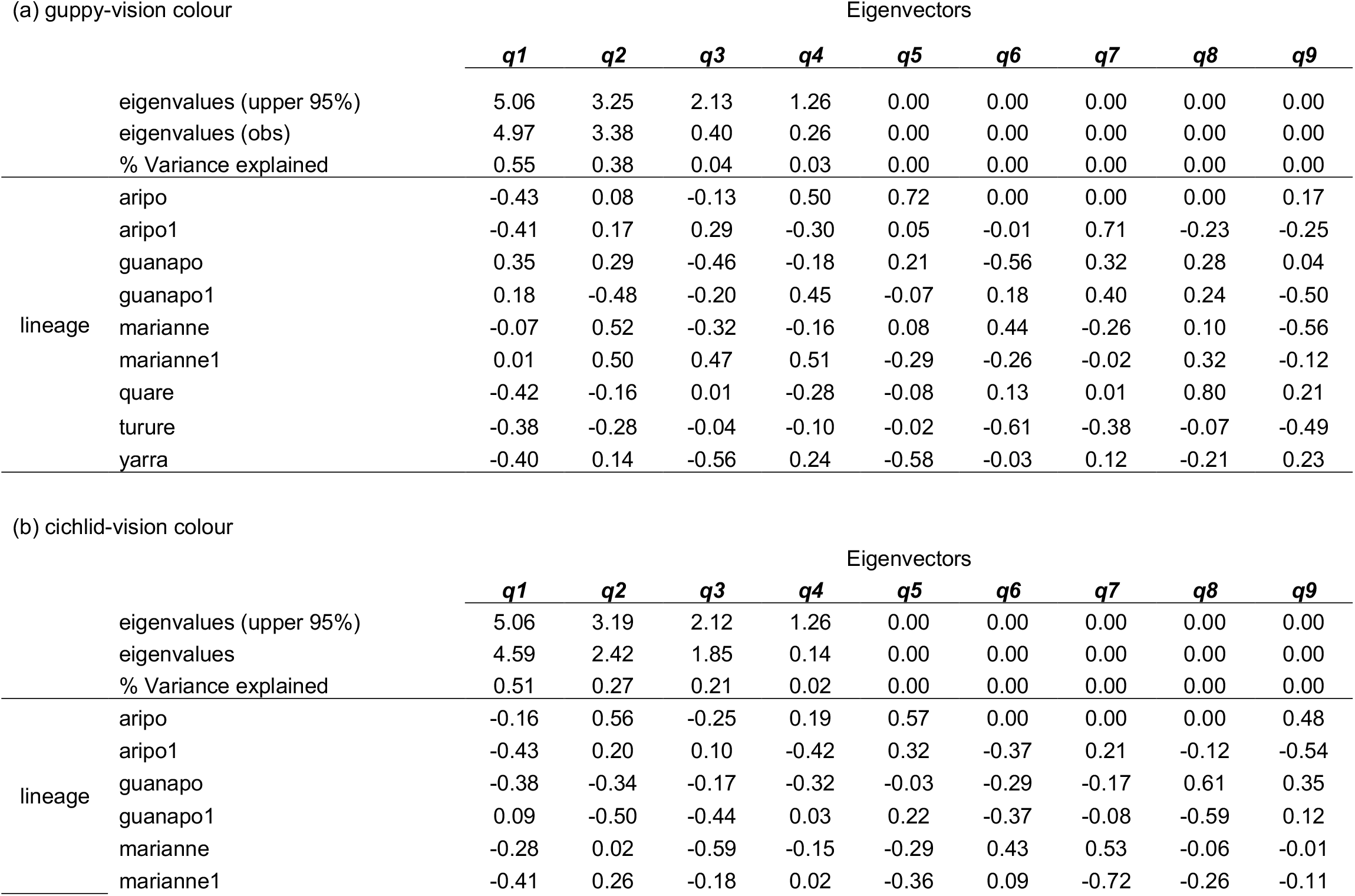
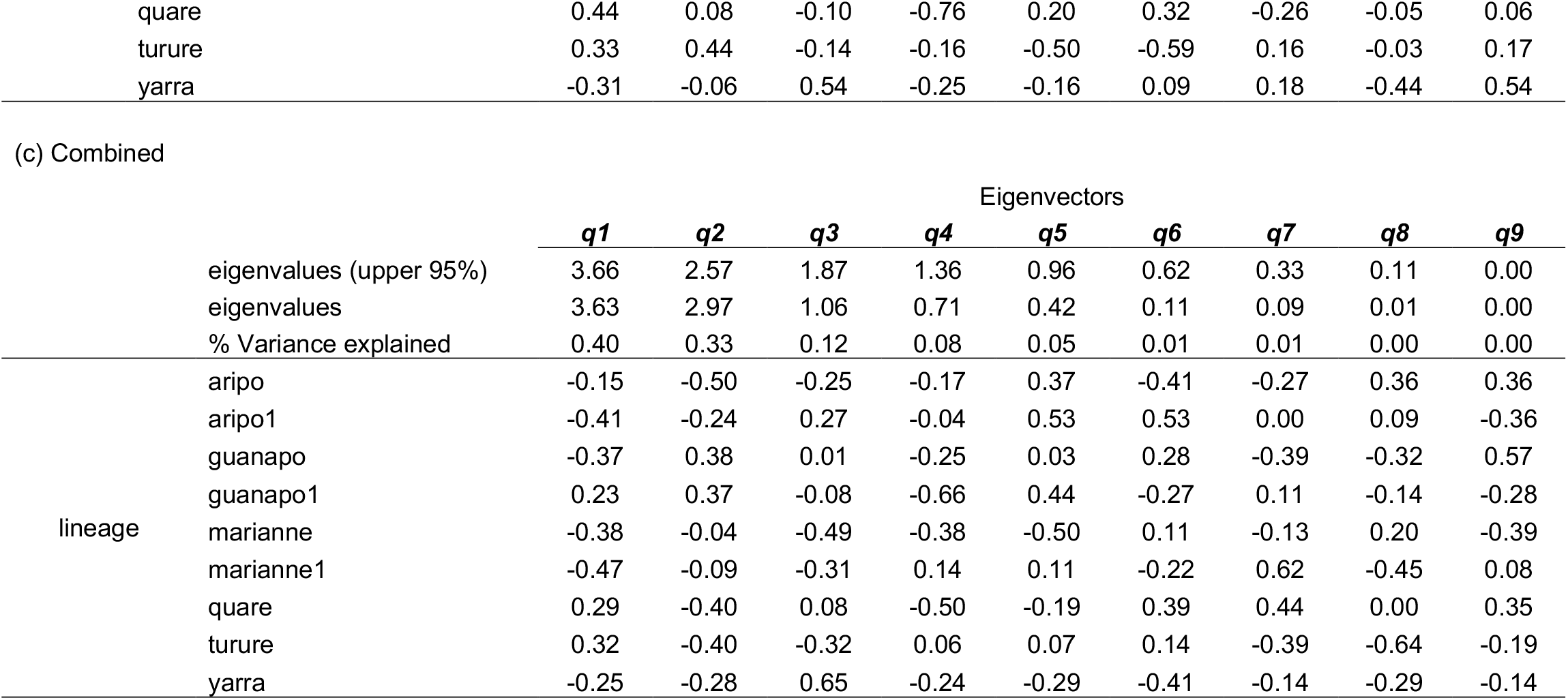
Spectral decomposition for among independent lineages for phenotypic parallelism. Note: Highlighted cells indicates that that dimension is significantly above 95% threshold.

### V. Genome-phenotype association analysis reveals that differentiated SNPs are found in genes related to cell morphogenesis and neural development

The core BayPass model revealed 85 SNPs more differentiated among populations than expected under neutrality at the 0.01% POD significance threshold (Figure 6). Of these, 61 and 24 were located within genic and non-genic regions, respectively (summarized in Table 5). Some of the genes were orthologous to zebrafish genes documented to be involved in cell morphogenesis and cell projections, colour patterning and pigmentation (e.g. *xpc, rpl, rilp, netrins*). Some were also within genes involved in neural development, such as sensory axon guidance (e.g. *ptprfa*). Using orthologous zebrafish genes determined by blasting, our GO analysis found no enrichment for any gene ontology category.

**Table 5.**
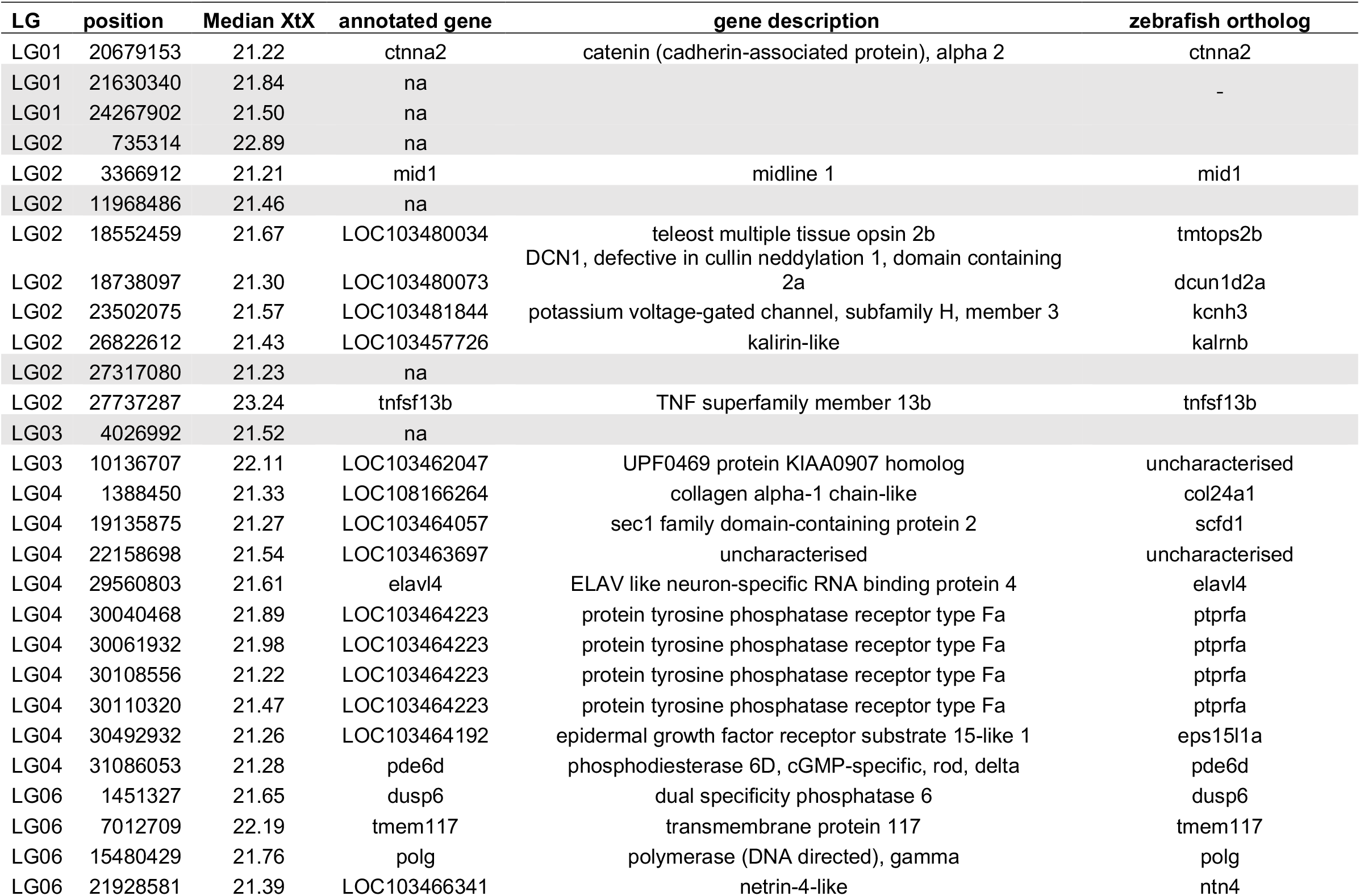
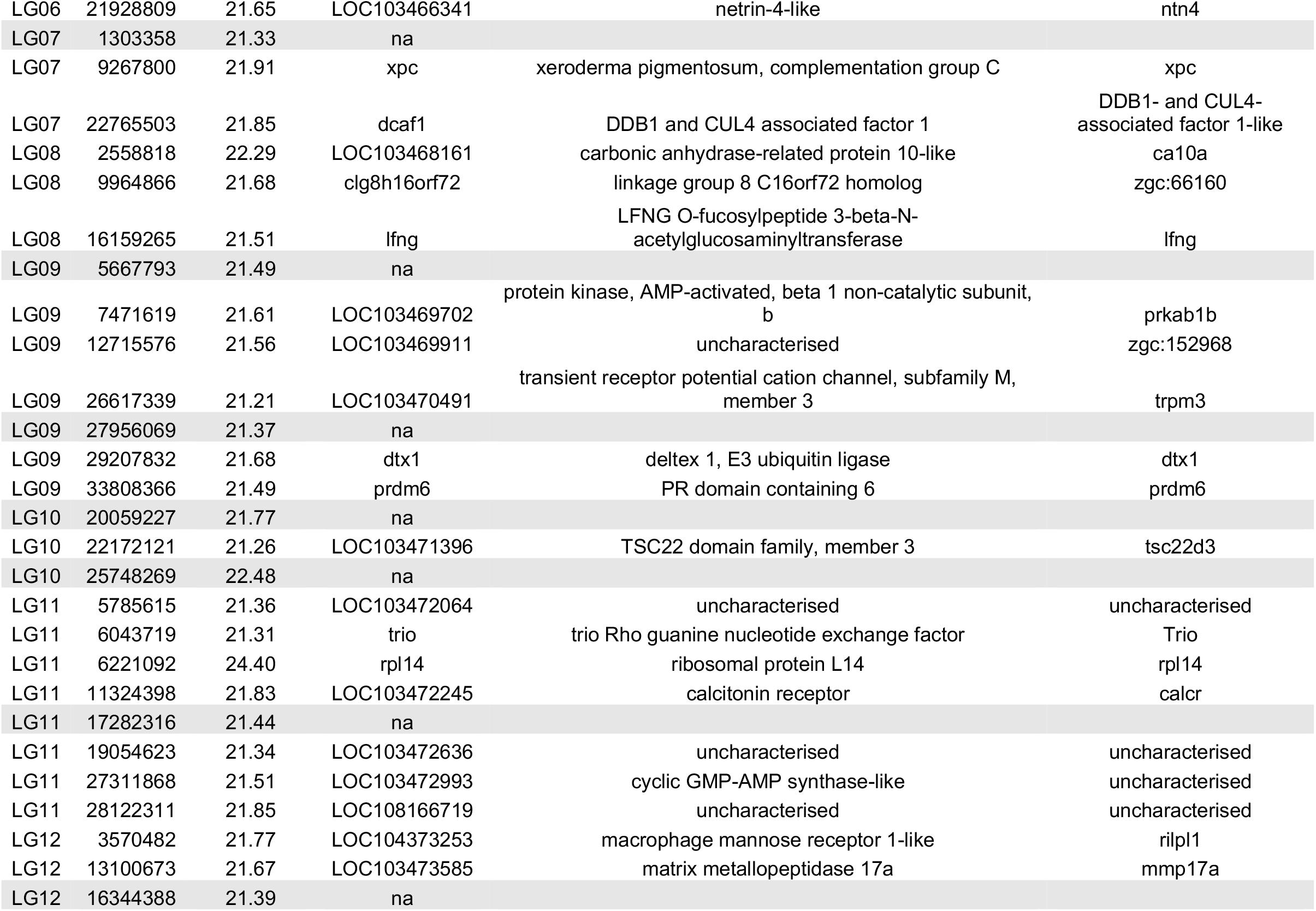
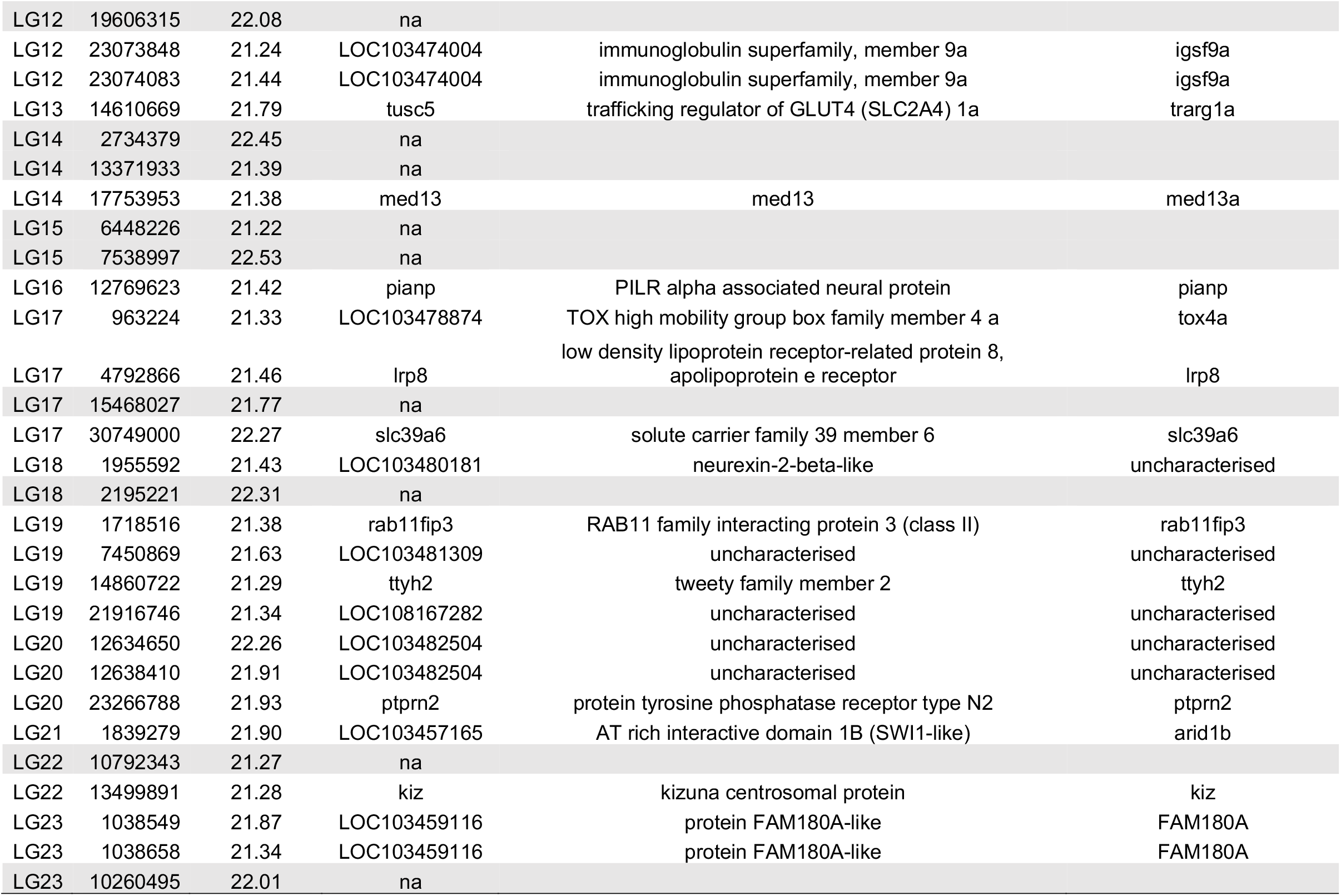
Location (LG and bp position) of significantly differentiated SNPs among populations and putative underlying genes. Shaded cells indicate non-genic regions within which SNPs are found.

**Figure 6.**
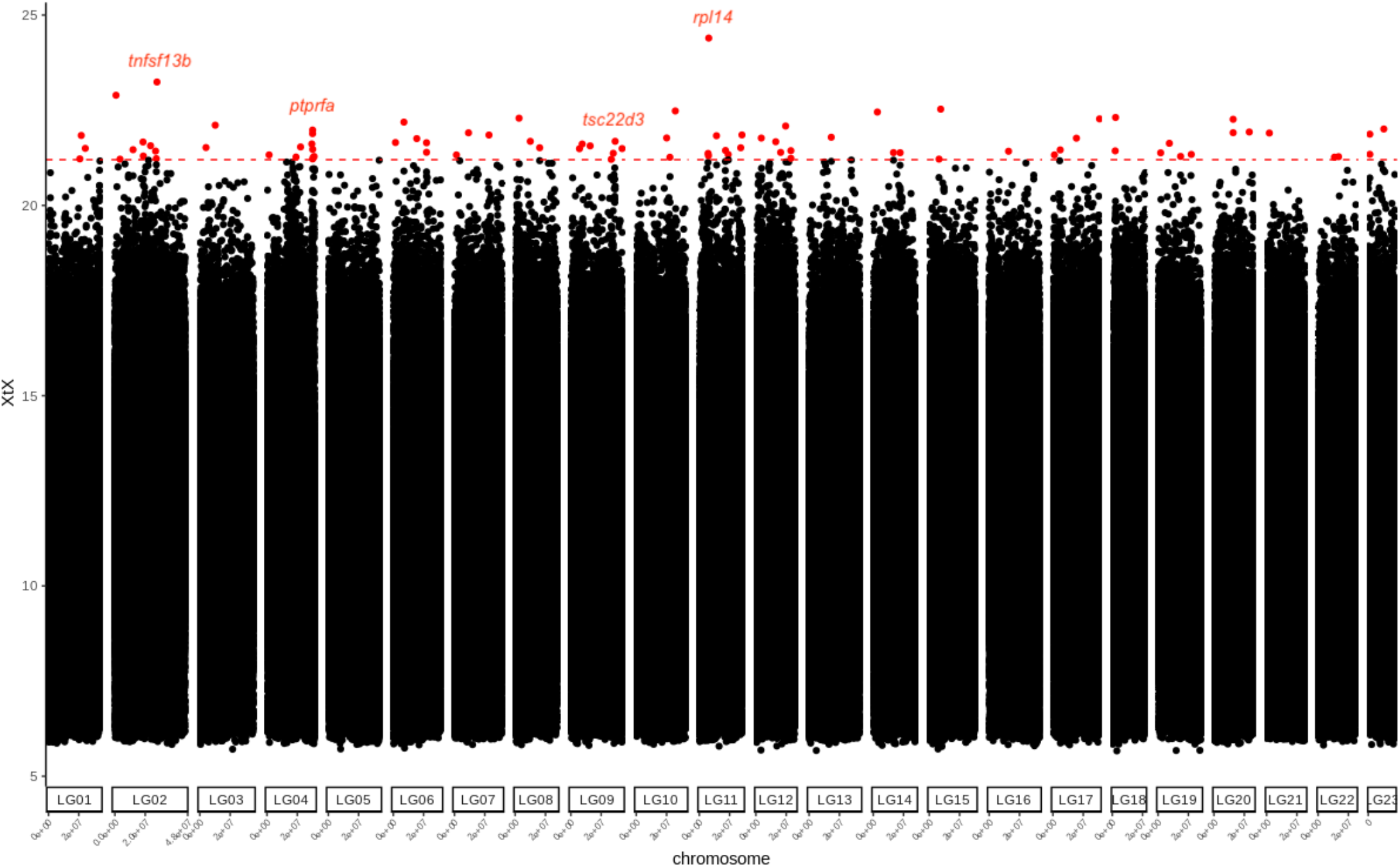
Manhattan plot of genome-wide differentiation among 12 male populations of guppies. The red dotted line and points indicate the 0.01% (21.2 *XtX*) threshold and significantly differentiated SNPs, respectively. Red dots denote significantly differentiated SNPs. Labels above dots denote putative candidate genes implicated cell morphogenesis linked to color patterns.

Using the BayPass IS model, we found a total of 195 SNPs associated with a tested covariate using a BF threshold > 30db (colour traits: guppy vision model = 77; cichlid vision model = 101; predation = 17, summarized in Table S7). Interestingly, there was no overlap with the 85 differentiated SNPs identified under the core model. Taken at face value, this suggests SNPs identified by the core model as being more differentiated than expected are not related to colour pattern differences among populations. SNPs significantly associated with covariates in the IS model were scattered throughout the genome (not clustered on specific chromosomes), and found within both non-genic and genic regions (∼42% in non-genic regions). With a single exception (LG22: 9903588; associated with both measures of sat_ΔS_) SNP-colour trait associations differed between between the guppy and cichlid visual perceptual models, consistent with the finding that traits defined under the two models have different genetic underpinnings. As with the *XtX*-based results from the core model, many SNPs associated with covariates under the IS model were found in genes previously implicated with cell morphogenesis and neuronal functions (e.g., *hapln2*; Table S7). There was no enrichment of genes related to any functional processes using putative genes identified from SNPs associated with traits under the guppy visual model. However, we found enrichment of genes for nervous system development, anatomical structure morphogenesis, and cellular process using SNPs associated with the traits defined usng the cichlid visual model (P < 0.0001, Table S8). Interestingly, some of the non-genic SNPs with elevated BF values were near genes that have known roles in teleost patterning (e.g. *bnc2, cdh11*, for full list see Table S9 and S10).

## DISCUSSION

Using colour traits defined by the visual systems of two major agents of selection on colour in male guppies (conspecifics, cichlid predator), we found an overall tendency towards higher, more conspicuous, phenotypic means in LP guppies as predicted. However, (i) trait means are not systematically higher at LP for within-river comparisons, (ii) statistical support for predation regime effects are weak, and (iii) regime effects explain little of the total among-population differentiation. Moreover, when modelling colour variation in multivariate phenotypic space we find (iv) little support for clustering of populations by predation regime, and (v) no evidence for parallel evolution of lineages. Nevertheless, putatively neutral patterns of genome-wide molecular differentiation did not readily explain the among-population phenotypic structure. This suggests that adaptive evolutionary processes have caused divergence of male colour among populations. Here, we first discuss these phenotypic patterns and their interpretation in relation to the evidence for parallel evolution of brighter male colouration across the high to low predation ecological transition in guppies. We then comment on the results of our genomic analyses that revealed some SNPs differentiated among populations in (or close) genes implicated in pigmentation, patterning and neuronal development in other fish species.

### Patterns of among-population phenotypic differentiation

Traits derived from the novel QCPA phenotyping pipeline offer only qualified support for the longstanding view that guppies from LP populations are particularly colourful and conspicuous (Haskins 1961; Endler 1980; Millar *et al*. 2006). Thus, trait means are higher at LP overall, but this is not a statistically robust pattern. Nor is it found consistently across the HP to LP transition within rivers. The lack of stronger patterns is perhaps somewhat surprising, although we note that previous guppy studies have also reported similarly inconsistent differences across regimes within rivers using transplant (Dick *et al*. 2018; Kemp *et al*. 2009, 2018) and predator-manipulation experiments (Gotanda *et al*. 2018). Colour differentiation between populations is strong using the QCPA phenotyping approach (approximately half of all phenotypic variance is among-populations), but predation regime does not explain much among-population variance (in single traits or multivariate phenotype). This conclusion is further supported by the finding that populations do not obviously cluster by predation regime in multi-trait space, regardless of whether this space is defined in 8 dimensions (using all traits defined from guppy and cichlid visual models) or 4 (i.e. using just guppy or just cichlid perception). Nor does correlation structure in the population variance-covariance matrix (**D)** supports the presence of a single major axis of variation running from low (i.e. low trait values) to high conspicuity.

In principle these results could occur even with parallel evolution of replicate lineages across the HP-LP transition, if recent coancestry, drift and/or ongoing gene flow between populations are masking the phenotypic signature of parallelism. We think multiple lines of evidence argue against this possibility. First, among the 12 populations with SNP data we found no correlation of pairwise genome wide F_ST_ values and phenotypic distance. Second, our use of mixed models to partition among-population variance suggest not only that neutral processes are insufficient to explain among-population differentiation, but also that adaptive divergence explains most variation (discussed further below). Third, and perhaps most importantly, when isolating the phenotypic change from HP to LP within rivers, the inferred directions of phenotypic evolution are not parallel among lineages. This analysis assumes that river can be used as a proxy for lineage (*sensu* de Lisle and Bolnick 2020), but that appears reasonable based on present and previous population genetic analyses (Willing *et al*. 2010; Blondel *et al*. 2020).

Thus, viewing male colour as a complex multivariate phenotype seen through the vision systems of biological agents of selection, we find no quantitative support for parallel evolution across the HP to LP regime transition. Rather, our results suggest that – in this phenotypic space – colour patterns evolve approximately independently in each river. However, the fact that genome wide molecular genetic structure does not predict phenotypic structuring argues against the idea that divergence of colour patterns could be primarily neutral. How can these results be reconciled with our existing understanding of colour evolution in guppies? In fact, we suggest the lack of parallelism, coupled to apparently greater variation among LP than among HP populations, is consistent with the widely-held view that reduced selection pressure from visual predators facilitates evolutionary divergence of male traits by female choice. In guppies, the evidence that females prefer more conspicuousness male patterns that also increase predation risk is abundant (reviewed in Houde 1997). However, this (univariate) conceptualisation of female choice masks the fact that, in multi-trait space, there may be many different directions that increase conspicuity to females.

The extent to which female choice drives Fisherian runaway evolution of traits that are ‘arbitrary’ (not under viability selection) versus ‘costly’ has been extensively debated (Kokko *et al*. 2002). However, even if females do prefer costly male phenotypes in all populations, this need not translate into closely aligned vectors of sexual selection on multivariate phenotype. Indeed, Endler and Houde (1995) demonstrated substantial geographic variation in female preferences for colour patches, e.g. black, orange, and colour contrast. Among-individual (female) differences in preference, a prerequisite for heritable variation that would allow trait-preference coevolution, have also been demonstrated (Brooks and Endler 2001; Brooks 2002). Lastly, we note that frequency dependent selection, with females preferring novel male phenotypes is well documented in guppies (Hughes *et al*. 2013), and will generally impose vectors of directional selection that differ in space (i.e. across populations) as well as time (e.g. as rare phenotypes become more common).

Our conclusion that predation regime has no consistent effect on the direction of evolution in multivariate trait holds irrespective of the vision model used to define phenotypes. However, the finding of population level correlations significantly less than +1 between homologous traits illustrates the wider potential for QPCA to offer new insights. The same objective colour phenotype may be perceived differently by prospective mates and potential predators. Here, guppy colour patterns appear generally more conspicuous in the chromatic channel for conspecifics, whereas the pike cichlid is more sensitive to achromatic (luminance) elements (Weadick *et al*. 2018). Such differences highlight the value of adopting colour measures appropriate to the visual systems of hypothesised selective agents (Endler 1978; Endler *et al*. 2005; Endler and Mielke, 2005), and have implications for the way we the evolution of colour signals with multiple receiving species. We also acknowledge however, that that our study is naive to any abiotic factors (e.g., substrate, water colour, light transmission, and canopy cover) that may affect *in situ* perception of phenotype by guppies and/or their predators. Local abiotic conditions are an explicit part of the wider sensory drive hypothesis (Endler 1980, 1992), and Kemp *et al*. (2018) found that canopy cover influenced the relative abundance of iridescent versus melanic colour patches in guppies. While we have no evidence to suggest this, it is at least possible that conditioning among-population (or lineage) variance on abiotic covariates would reveal greater support for parallel evolution in the colour phenotype as measured here.

### Insights from genome-scans and SNP associations

Our pool-sequencing data provide several genetic insights that complement the phenotypic analyses. First, based on detected associations with the phenotypic traits defined here, male colour patterns are probably highly polygenic with genes distributed across the genome rather than being restricted to the sex chromosomes or otherwise clustered. Associated SNPs are found both outside and within genic regions, and involve cis-acting elements. For instance, a SNP (LG:31463600) associated with differences in guppy-specific saturation was found in an intergenic region near basonuclin 2 (*bnc2*), a gene implicated in the maintenance of extracellular environments within which pigment cells driving patterning reside (Lang *et al*. 2009). Third, the minimal overlap between SNPs associated with homologous traits defined under the two vision models is consistent with the population level phenotypic correlations being less that one, suggesting that homolgous traits defined under different models are genetically distinct. However, since colour genetics have often been studied using human visual perception (Tripathi *et al*. 2009), it also highlights the possibility that ecologically-important genes may have been previously overlooked. Conversely, we did not detect any trait-SNP association consistency with colour genes previously identified in guppies, i.e. *csf1ra* and *kita* (Kottler *et al*. 2013). This may again be explained by our choice of defined phenotype; focusing here on measures of overall body patterns and/or conspicuity (as perceived by selective agents), rather than features of individual pigmented patches (as perceived by humans). Fourth, no SNPs significantly associated with tested covariables were actually among the set identified as being significantly more differentiated than expected under neutrality in the core BayPass model. Taken at face value, this could indicate that greater (adaptive) genetic divergence among populations has occurred for specific traits or aspects of phenotype (e.g. life history) that were not quantified in this study.

To our knowledge, this is the first study to investigate among-population natural variation in male guppy patterns at the genomic level. Consequently some brief, and necessarily tentative, comments on specific SNPs identified are perhaps warranted. Among those more differentiated than expected under neutrality, several SNPs were detected in the protein tyrosine phosphatase receptor of types *Fa* (*ptprfa*) and N2 (*ptprn2*). Basic functions of these genes include cell proliferation and epithelial cell-cell adhesion. They are considered to counteract tyrosinase kinase receptors (Xu and Fisher, 2012), which play known roles in melanocyte and melanophores development (*Kita* and *cKit*, respectively; Alexeev and Yoon 2006; Larsson and Parichy 2019). It is not known if tyrosine phosphatase receptors carry a direct pigmentation function, although this is plausible given their documented involvement in cell differentiation, oncogenic-events, and sensory guidance to the skin (Wang *et al*.2012). Several SNPs deviating from neutral expectaions were also identified in less well-known genes including *polg* and *trio*, which are part of regulatory networks influencing melanocyte differentiation (Seberg *et al*. 2017; Park *et al*. 2018). Among SNPs associated with tested covariates (trait means and predation regime), we also find several in (or near) genes with known roles in fish colour. For instance, one SNP association with guppy vision sat_ΔS_ was near *ablim3*, a gene thought to be involved in pigment cell movement in fishes (Ahi *et al*. 2020). Another was found within *cx30*.*3*, part of a family of genes implicated in zebrafish skin development and pattern formation (Tao *et al*. 2010; Irion *et al*. 2014).

## Conclusion

Trinidadian guppies ostensibly represent a classic example of parallel (or convergent) evolution, with repeated divergence in male colour patterns between upstream and downstream populations due partly to predation conditions. We find that the well-described tendency for greater conspicuousness under low predation holds qualitatively true when we analyse single traits defined under guppy and predator vision models. However, statistical support for this is weak, and repeated colonisation of low predation habitats has not lead to the parallel evolution of conspicuous phenotypes when these are characterised in quantitative, multivariate, phenotypic space. Instead, we suggest that colonisation of LP habitat may reduce selective constraints on phenotypic space imposed by predation, facilitating population-specific divergence under sexual selection. Together, our results are nonetheless consistent with the standard view that reduced selection by predators allows male colour traits to evolve under sexual selection imposed by female choice. However, the common assertion that female choice in guppies selects for ‘brighter’ or more ‘conspicuous’ males, masks the fact that – in multivariate trait space - colour is evolving in rather different directions across populations and lineages.

## ACKNOWLEDGEMENTS

This work was supported by a BBSRC grant to AJW (BB/L022656/1), a Genetics Society (UK) Grant to LY and a European Research Council Advanced Grant 695225 (GUPPYSEX) awarded to D. Charlesworth (with AJW and DC as co-investigators). We thank D. Charlesworth for comments on an earlier version of this MS. We also thank R. Mahabir, R. Heathcote for assistance with the field collection and Trinidad-UK shipment of guppies. Guppy collection was approved by Trinidad’s Ministry of Agriculture, Land and Fisheries and import to the UK was approved by CEFAS (EW087-O-816A).

## DATA ACCESSIBILITY

Raw phenotypic and genomic data as well as R and Java scripts will be available publicly upon the acceptance of the manuscript in Dryad, GeneBank, and appropriate data repositories.

## AUTHOR CONTRIBUTIONS

LY and AW designed the study with input from DPC and JT. LY, DPC and AW performed the research. JT and IR contributed analytical tools. LY and AW analysed the data. LY and AW wrote the paper with input from DCP, IR, and JT.

## Figure Legends

**Figure S1.**
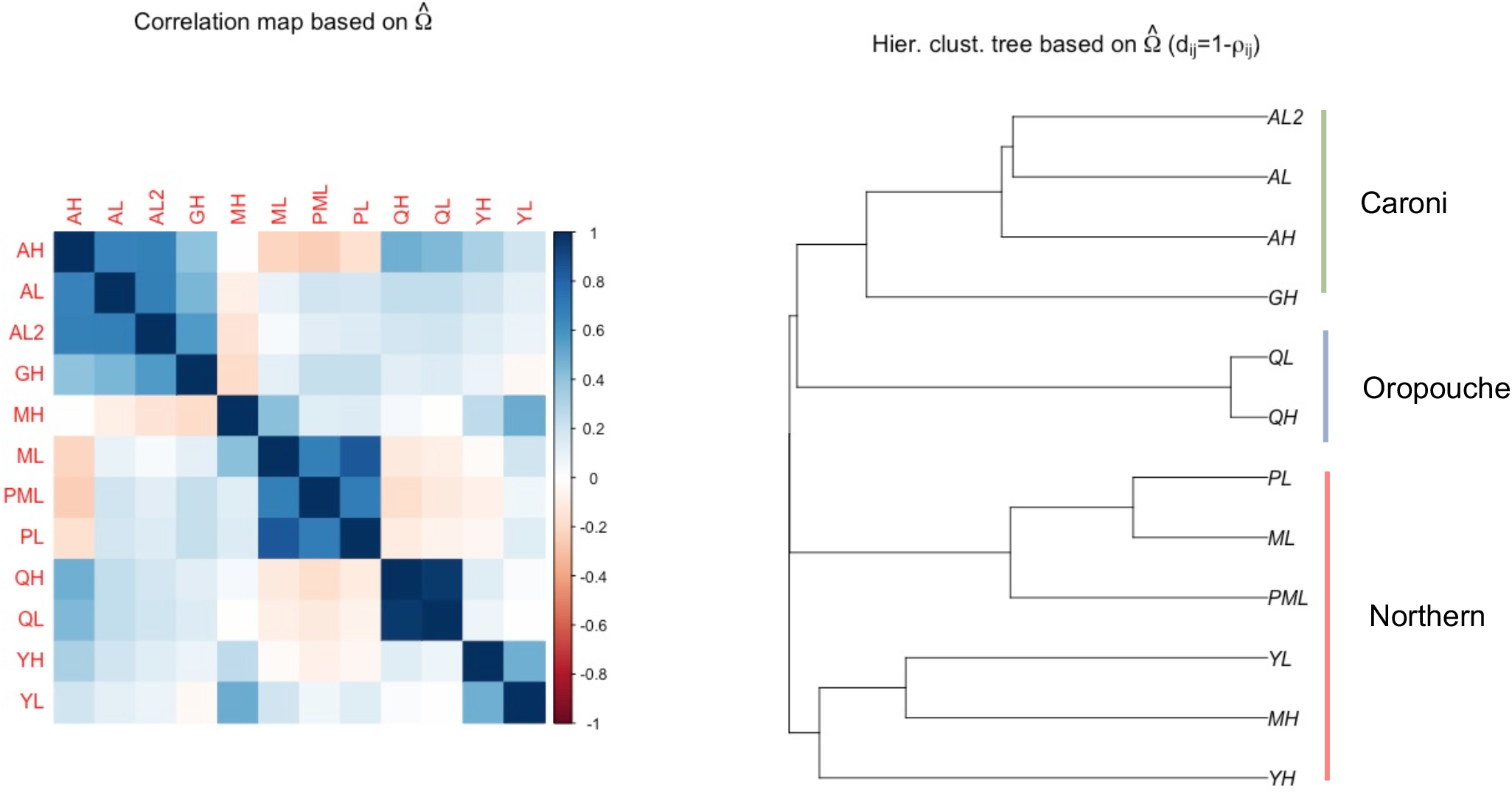
Illustration of the covariance matrix **Ω** among 12 populations estimated from BayPass core model using ca. 100,000 SNPs.

**Figure S2.**
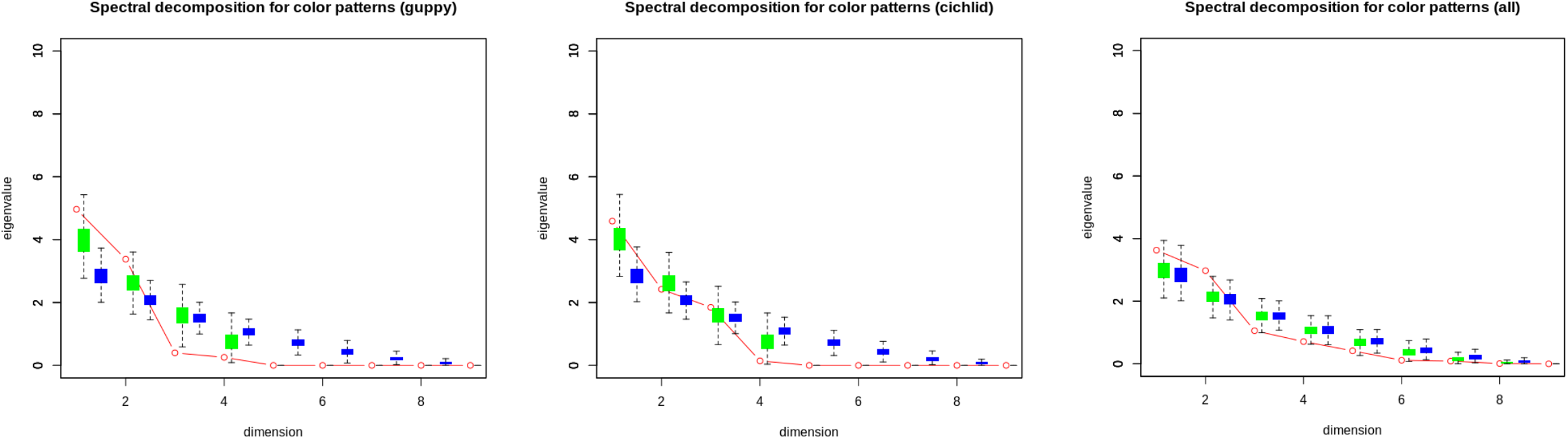
Eigen decomposition of C matrices for (a) guppy-(b) cichlid-(c) guppy and cichlid-perceived color patterns, demonstrating general support for the null hypothesis of independent change in color patterns. Red dots and line indicate observed values from diagonalization of the estimated matrix C. Per De Lisle and Bolnick (2020), green and boxplots represent the expected distribution of eigenvalues under a null hypothesis of a random direction of evolutionary change across lineages, calculated from sampling the corresponding Wishart distribution or an simulated distribution by placing random vectors in a trait space (1000 simulations)

## REFERENCES

Ahi, E. P., Lecaudey, L. A., Ziegelbecker, A., Steiner, O., Glabonjat, R., Goessler, W., … Sefc, K. M. (2020). Comparative transcriptomics reveals candidate carotenoid color genes in an East African cichlid fish. BMC Genomics, 21. doi: 10.1186/s12864-020-6473-8

Alexeev, V., & Yoon, K. (2006). Distinctive role of the cKit receptor tyrosine kinase signaling in mammalian melanocytes. The Journal of Investigative Dermatology, 126(5), 1102– 1110. doi: 10.1038/sj.jid.5700125

Allender, C. J., Seehausen, O., Knight, M. E., Turner, G. F., & Maclean, N. (2003). Divergent selection during speciation of Lake Malawi cichlid fishes inferred from parallel radiations in nuptial coloration. Proceedings of the National Academy of Sciences, 100(24), 14074–14079. doi: 10.1073/pnas.2332665100

Ashburner, M., Ball, C. A., Blake, J. A., Botstein, D., Butler, H., Cherry, J. M., … Sherlock, G. (2000). Gene ontology: tool for the unification of biology. The Gene Ontology Consortium. Nature Genetics, 25(1), 25–29. doi: 10.1038/75556

Astles, P. A., Moore A. J., & Preziosi R. F. (2006). A comparison of methods to estimate cross-environment genetic correlations. Journal of Evolutionary Biology 19(1): 114–122. doi: 10.1111/j.1420-9101.2005.00997.x.

Belleghem, S. M. V., Papa, R., Ortiz-Zuazaga, H., Hendrickx, F., Jiggins, C. D., McMillan, W. O., & Counterman, B. A. (2018). patternize: An R package for quantifying colour pattern variation. Methods in Ecology and Evolution, 9(2), 390–398. doi: 10.1111/2041-210X.12853

Blondel, L., Baillie, L., Quinton, J., Alemu, J. B., Paterson, I., Hendry, A. P., & Bentzen, P. (2019). Evidence for contemporary and historical gene flow between guppy populations in different watersheds, with a test for associations with adaptive traits. Ecology and Evolution, 9(8), 4504–4517. doi: 10.1002/ece3.5033

Bolnick, D. I., Barrett, R. D. H., Oke, K. B., Rennison, D. J., & Stuart, Y. E. (2018). (Non)Parallel Evolution. Annual Review of Ecology, Evolution, and Systematics, 49(1), 303–330. doi: 10.1146/annurev-ecolsys-110617-062240

Brooks, R., & Endler, J. A. (2001). Female guppies agree to differ: phenotypic and genetic variation in mate-choice behavior and the consequences for sexual selection. Evolution 55(8), 1644–1655. doi: 10.1111/j.0014-3820.2001.tb00684.x

Brooks, Robert. (2002). Variation in female mate choice within guppy populations: population divergence, multiple ornaments and the maintenance of polymorphism. Genetica, 116(2–3), 343–358.

Butler, D., Cullis, B., Gilmour, A., Gogel, B., & Thompson, R. (2017). ASReml-R Reference Manual Version 4. Hemel Hempstead, HP1 1ES, UK: VSN International Ltd.

Butlin, R. K., Saura, M., Charrier, G., Jackson, B., André, C., Caballero, A., … Rolán-Alvarez, E. (2014). Parallel evolution of local adaptation and reproductive isolation in the face of gene flow. Evolution; International Journal of Organic Evolution, 68(4), 935–949. doi: 10.1111/evo.12329

Caves, E. M., Sutton, T. T., & Johnsen, S. (2017). Visual acuity in ray-finned fishes correlates with eye size and habitat. The Journal of Experimental Biology, 220(Pt 9), 1586–1596. doi: 10.1242/jeb.151183

Cole, G. L., & Endler, J. A. (2015). Variable environmental effects on a multicomponent sexually selected trait. The American Naturalist, 185(4), 452–468. doi: 10.1086/680022

Cuthill, I. C., Allen, W. L., Arbuckle, K., Caspers, B., Chaplin, G., Hauber, M. E., … Caro, T. (2017). The biology of color. Science, 357(6350). doi: 10.1126/science.aan0221

De Lisle, S. P. D., & Bolnick, D. I. (2020). A multivariate view of parallel evolution. Evolution, 74(7), 1466–1481. doi: 10.1111/evo.14035

Dick, C., Hinh, J., Hayashi, C. Y., & Reznick, D. N. (2018). Convergent evolution of coloration in experimental introductions of the guppy (Poecilia reticulata). Ecology and Evolution, 8(17), 8999–9006. doi: 10.1002/ece3.4418

Dyer, A. G., Boyd-Gerny, S., McLoughlin, S., Rosa, M. G. P., Simonov, V., & Wong, B. B. M. (2012). Parallel evolution of angiosperm colour signals: common evolutionary pressures linked to hymenopteran vision. Proceedings. Biological Sciences, 279(1742), 3606–3615. doi: 10.1098/rspb.2012.0827

Elmer, K. R., & Meyer, A. (2011). Adaptation in the age of ecological genomics: insights from parallelism and convergence. Trends in Ecology & Evolution, 26(6), 298–306. doi: 10.1016/j.tree.2011.02.008

Endler, J. A. (1978). A Predator’s View of Animal Color Patterns. Evolutionary Biology, 319– 364. doi: 10.1007/978-1-4615-6956-5_5

Endler, J. A. (1980). Natural Selection on Color Patterns in Poecilia reticulata. Evolution, 34(1), 76–91. JSTOR. doi: 10.2307/2408316

Endler, J. A. (1983). Natural and secual selection on color patterns in poeciliids fishers. Environment Biology of Fishes 9(2), 173–190.

Endler, J. A. (1991). Variation in the appearance of guppy color patterns to guppies and their predators under different visual conditions. Vision Research, 31(3), 587–608. doi: 10.1016/0042-6989(91)90109-i

Endler, J. A. (1992). Signals, Signal Conditions, and the Direction of Evolution. The American Naturalist, 139, S125–S153. doi: 10.1086/285308

Endler, J. A. (1995). Multiple-trait coevolution and environmental gradients in guppies. Trends in Ecology & Evolution, 10(1), 22–29. doi: 10.1016/s0169-5347(00)88956-9

Endler, J. A. (2012). A framework for analysing colour pattern geometry: adjacent colours. Biological Journal of the Linnean Society, 107(2), 233–253. doi: 10.1111/j.1095-8312.2012.01937.x

Endler, J. A. (2015). Signals, Signal Conditions, and the Direction of Evolution. The American Naturalist. (world). doi: 10.1086/285308

Endler, J. A., Cole, G. L., & Kranz, A. M. (2018). Boundary strength analysis: Combining colour pattern geometry and coloured patch visual properties for use in predicting behaviour and fitness. Methods in Ecology and Evolution, 9(12), 2334–2348. doi: 10.1111/2041-210X.13073

Endler, J. A., & Houde, A. E. (1995). Geographic variation in female preferences for male traits in poecilia reticulata. Evolution, 49(3), 456–468. doi: 10.1111/j.1558-5646.1995.tb02278.x

Endler, J. A., & Mappes, J. (2017). The current and future state of animal coloration research. Philosophical Transactions of the Royal Society of London. Series B, Biological Sciences, 372(1724). doi: 10.1098/rstb.2016.0352

Endler, J. A., & Mielke, P. W. (2005). Comparing entire colour patterns as birds see them. Biological Journal of the Linnean Society, 86(4), 405–431. doi: 10.1111/j.1095-8312.2005.00540.x

Fraser, B. A., Künstner, A., Reznick, D. N., Dreyer, C., & Weigel, D. (2015). Population genomics of natural and experimental populations of guppies (Poecilia reticulata). Molecular Ecology, 24(2), 389–408. doi: 10.1111/mec.13022

Funk, D. J., Egan, S. P., & Nosil, P. (2011). Isolation by adaptation in Neochlamisus leaf beetles: host-related selection promotes neutral genomic divergence. Molecular Ecology, 20(22), 4671–4682. doi: 10.1111/j.1365-294X.2011.05311.x

Gautier, M. (2015). Genome-Wide Scan for Adaptive Divergence and Association with Population-Specific Covariates. Genetics, 201(4), 1555–1579. doi: 10.1534/genetics.115.181453

Gautier, M., Foucaud, J., Gharbi, K., Cézard, T., Galan, M., Loiseau, A., … Estoup, A. (2013). Estimation of population allele frequencies from next-generation sequencing data: pool-versus individual-based genotyping. Molecular Ecology, 22(14), 3766– 3779. doi: 10.1111/mec.12360

Gautier, M., Yamaguchi, J., Foucaud, J., Loiseau, A., Ausset, A., Facon, B., … Prud’homme, B. (2018). The Genomic Basis of Color Pattern Polymorphism in the Harlequin Ladybird. Current Biology: CB, 28(20), 3296-3302.e7. doi: 10.1016/j.cub.2018.08.023

Gotanda, K. M., Pack, A., LeBlond, C., & Hendry, A. P. (2019). Do replicates of independent guppy lineages evolve similarly in a predator-free laboratory environment? Ecology and Evolution, 9(1), 36–51. doi: 10.1002/ece3.4585

Günther, T., & Coop, G. (2013). Robust identification of local adaptation from allele frequencies. Genetics, 195(1), 205–220. doi: 10.1534/genetics.113.152462

Hartl, D. L., & Clark, A. G. (1997). Principles of population genetics. Sinauer Associates.

Haskins, C. P., Haskins, E. F., McLaughlin, J. J., & Hewitt, R. E. (1961). Polymorphism and population structure in Lebistes reticulatus, an ecological study. In Vertebrate Speciation (W. F. Blair, pp. 320–395). Austin: University of Texas Press.

Houde, A. (1997). Sex, Color, and Mate Choice in Guppies (Vol. 71). Princeton University Press. JSTOR. doi: 10.2307/j.ctvs32rtk

Howe, K., Clark, M. D., Torroja, C. F., Torrance, J., Berthelot, C., Muffato, M., … Stemple, D. L. (2013). The zebrafish reference genome sequence and its relationship to the human genome. Nature, 496(7446), 498–503. doi: 10.1038/nature12111

Hughes, K. A., Houde, A. E., Price, A. C., & Rodd, F. H. (2013). Mating advantage for rare males in wild guppy populations. Nature, 503(7474), 108–110. doi: 10.1038/nature12717

Irion, U., Frohnhöfer, H. G., Krauss, J., Champollion, T. Ç., Maischein, H.-M., Geiger-Rudolph, S., … Nüsslein-Volhard, C. (2014, December 23). Gap junctions composed of connexins 41.8 and 39.4 are essential for colour pattern formation in zebrafish. doi: 10.7554/eLife.05125

Karim, N., Gordon, S. P., Schwartz, A. K., & Hendry, A. P. (2007). This is not déjà vu all over again: male guppy colour in a new experimental introduction. Journal of Evolutionary Biology, 20(4), 1339–1350. doi: 10.1111/j.1420-9101.2007.01350.x

Kawamura, S., Kasagi, S., Kasai, D., Tezuka, A., Shoji, A., Takahashi, A., … Kawata, M. (2016). Spectral sensitivity of guppy visual pigments reconstituted in vitro to resolve association of opsins with cone cell types. Vision Research, 127, 67–73. doi: 10.1016/j.visres.2016.06.013

Kemp, D. J., Batistic, F.-K., & Reznick, D. N. (2018). Predictable adaptive trajectories of sexual coloration in the wild: Evidence from replicate experimental guppy populations*. Evolution, 72(11), 2462–2477. doi: 10.1111/evo.13564

Kemp, D. J., Reznick, D. N., Grether, G. F., & Endler, J. A. (2009). Predicting the direction of ornament evolution in Trinidadian guppies (Poecilia reticulata). Proceedings of the Royal Society B: Biological Sciences. (world). doi: 10.1098/rspb.2009.1226

Kofler, R., Pandey, R. V., & Schlötterer, C. (2011). PoPoolation2: identifying differentiation between populations using sequencing of pooled DNA samples (Pool-Seq). Bioinformatics, 27(24), 3435–3436. doi: 10.1093/bioinformatics/btr589

Kokko, H., Brooks, R., McNamara, J. M., & Houston, A. I. (2002). The sexual selection continuum. Proceedings of the Royal Society of London. Series B: Biological Sciences. (world). doi: 10.1098/rspb.2002.2020

Kottler, V. A., Fadeev, A., Weigel, D., & Dreyer, C. (2013). Pigment pattern formation in the guppy, Poecilia reticulata, involves the Kita and Csf1ra receptor tyrosine kinases. Genetics, 194(3), 631–646. doi: 10.1534/genetics.113.151738

Kranz, A. M., Cole, G. L., Singh, P., & Endler, J. A. (2018). Colour pattern component phenotypic divergence can be predicted by the light environment. Journal of Evolutionary Biology, 31(10), 1459–1476. doi: 10.1111/jeb.13342

Kranz, A. M., Forgan, L. G., Cole, G. L., & Endler, J. A. (2018). Light environment change induces differential expression of guppy opsins in a multi-generational evolution experiment. Evolution 72(8), 1656–1676. doi: 10.1111/evo.13519

Kratochwil, C. F., Liang, Y., Gerwin, J., Woltering, J. M., Urban, S., Henning, F., … Meyer, A. (2018). Agouti-related peptide 2 facilitates convergent evolution of stripe patterns across cichlid fish radiations. Science (New York, N.Y.), 362(6413), 457–460. doi: 10.1126/science.aao6809

Künstner, A., Hoffmann, M., Fraser, B. A., Kottler, V. A., Sharma, E., Weigel, D., & Dreyer, C. (2016). The Genome of the Trinidadian Guppy, Poecilia reticulata, and Variation in the Guanapo Population. PloS One, 11(12), e0169087. doi: 10.1371/journal.pone.0169087

Lang, M. R., Patterson, L. B., Gordon, T. N., Johnson, S. L., & Parichy, D. M. (2009). Basonuclin-2 Requirements for Zebrafish Adult Pigment Pattern Development and Female Fertility. PLOS Genetics, 5(11), e1000744. doi: 10.1371/journal.pgen.1000744

Laver, C. R. J., & Taylor, J. S. (2011). RT-qPCR reveals opsin gene upregulation associated with age and sex in guppies (Poecilia reticulata) - a species with color-based sexual selection and 11 visual-opsin genes. BMC Evolutionary Biology, 11, 81. doi: 10.1186/1471-2148-11-81

Leinonen, T., McCairns, R. J. S., O’Hara, R. B., & Merilä, J. (2013). Q ST – F ST comparisons: evolutionary and ecological insights from genomic heterogeneity. Nature Reviews Genetics, 14(3), 179–190. doi: 10.1038/nrg3395

Li, H., & Durbin, R. (2010). Fast and accurate long-read alignment with Burrows-Wheeler transform. Bioinformatics (Oxford, England), 26(5), 589–595. doi: 10.1093/bioinformatics/btp698

Li, H., Handsaker, B., Wysoker, A., Fennell, T., Ruan, J., Homer, N., … 1000 Genome Project Data Processing Subgroup. (2009). The Sequence Alignment/Map format and SAMtools. Bioinformatics (Oxford, England), 25(16), 2078–2079. doi: 10.1093/bioinformatics/btp352

Lindholm A, Breden F. (2002). Sex chromosomes and sexual selection in Poeciliid Fishes. American Naturalist 160, S214–224.

Long, K. D. (n.d.). Variation in mating behavior of the guppy, Poecilia reticulata, as a function of environmental, irradiance, visual acuity, and perception of male color patern (Doctoral dissertation). Berkeley CA: University of California.

Magurran, A. E. (n.d.). Evolutionary Ecology: The Trinidadian Guppy. Oxford University Press.

Martin, R. A., Riesch, R., Heinen-Kay, J. L., & Langerhans, R. B. (2014). Evolution of male coloration during a post-Pleistocene radiation of Bahamas mosquitofish (Gambusia hubbsi). Evolution, 68(2), 397–411. doi: 10.1111/evo.12277

McKinnon, J. S., Mori, S., Blackman, B. K., David, L., Kingsley, D. M., Jamieson, L., … Schluter, D. (2004). Evidence for ecology’s role in speciation. Nature, 429(6989), 294–298. doi: 10.1038/nature02556

Millar, N. P., Reznick, D. N., Kinnison, M. T., & Hendry, A. P. (2006). Disentangling the selective factors that act on male colour in wild guppies. Oikos, 113(1), 1–12. doi: 10.1111/j.0030-1299.2006.14038.x

Montenegro, J., Mochida, K., Matsui, K., Mokodongan, D. F., Sumarto, B. K. A., Lawelle, S. A., … Yamahira, K. (2019). Convergent evolution of body color between sympatric freshwater fishes via different visual sensory evolution. Ecology and Evolution, 9(11), 6389–6398. doi: 10.1002/ece3.5211

Nosil, P., Egan, S. P., & Funk, D. J. (2008). Heterogeneous genomic differentiation between walking-stick ecotypes: “isolation by adaptation” and multiple roles for divergent selection. Evolution; International Journal of Organic Evolution, 62(2), 316–336. doi: 10.1111/j.1558-5646.2007.00299.x

Nosil, P., Funk, D. J., & Ortiz-Barrientos, D. (2009). Divergent selection and heterogeneous genomic divergence. Molecular Ecology, 18(3), 375–402. doi: 10.1111/j.1365-294X.2008.03946.x

Olendorf, R., Rodd, F. H., Punzalan, D., Houde, A. E., Hurt, C., Reznick, D. N., & Hughes, K. A. (2006). Frequency-dependent survival in natural guppy populations. Nature, 441(7093), 633–636. doi: 10.1038/nature04646

Park, J. J., Diefenbach, R. J., Joshua, A. M., Kefford, R. F., Carlino, M. S., & Rizos, H. (2018). Oncogenic signaling in uveal melanoma. Pigment Cell & Melanoma Research, 31(6), 661–672. doi: 10.1111/pcmr.12708

Pascoal, S., Mendrok, M., Mitchell, C., Wilson, A. J., Hunt, J., & Bailey, N. W. (2016). Sexual selection and population divergence I: The influence of socially flexible cuticular hydrocarbon expression in male field crickets (Teleogryllus oceanicus). Evolution, 70(1), 82–97. doi: 10.1111/evo.12839

Pascoal, S., Mendrok, M., Wilson, A. J., Hunt, J., & Bailey, N. W. (2017). Sexual selection and population divergence II. Divergence in different sexual traits and signal modalities in field crickets (Teleogryllus oceanicus). Evolution, 71(6), 1614–1626. doi: 10.1111/evo.13239

Patterson, L. B., & Parichy, D. M. (2019). Zebrafish Pigment Pattern Formation: Insights into the Development and Evolution of Adult Form. Annual Review of Genetics, 53, 505– 530. doi: 10.1146/annurev-genet-112618-043741

Pujol, B., Wilson, A. J., Ross, R. I. C., & Pannell, J. R. (2008). Are QST–FST comparisons for natural populations meaningful? Molecular Ecology, 17(22), 4782–4785. doi: 10.1111/j.1365-294X.2008.03958.x

R Core Team. (2019). R: A language and environment for statistical computing. Retrieved June 10, 2020, from R Foundation for Statistical Computing, Vienna, Austria website: https://www.r-project.org/

Räsänen, K., & Hendry, A. P. (2008). Disentangling interactions between adaptive divergence and gene flow when ecology drives diversification. Ecology Letters, 11(6), 624–636. doi: 10.1111/j.1461-0248.2008.01176.x

Rasband, W. S. (1997). ImageJ. Bethesda, Maryland, USA: U. S. National Institutes of Health. Retrieved from https://imagej.nih.gov/ij/

Reznick, D. N. (1997). Life history evolution in guppies (Poecilia reticulata): guppies as a model for studying the evolutionary biology of aging. Experimental Gerontology, 32(3), 245–258. doi: 10.1016/s0531-5565(96)00129-5

Reznick, David N., Butler, M. J., Rodd, F. H., & Ross, P. (1996). Life-history evolution in guppies (Poecilia reticulata) 6. Differential mortality as a mechanism for natural selection. Evolution, 50(4), 1651–1660. doi: 10.1111/j.1558-5646.1996.tb03937.x

Schlötterer, C., Tobler, R., Kofler, R., & Nolte, V. (2014a). Sequencing pools of individuals - mining genome-wide polymorphism data without big funding. Nature Reviews. Genetics, 15(11), 749–763. doi: 10.1038/nrg3803

Schluter, D., Clifford, E. A., Nemethy, M., & McKinnon, J. S. (2004). Parallel evolution and inheritance of quantitative traits. The American Naturalist, 163(6), 809–822. doi: 10.1086/383621

Seberg, H. E., Van Otterloo, E., Loftus, S. K., Liu, H., Bonde, G., Sompallae, R., … Cornell, R. A. (2017). TFAP2 paralogs regulate melanocyte differentiation in parallel with MITF. PLoS Genetics, 13(3). doi: 10.1371/journal.pgen.1006636

Seehausen, O., Terai, Y., Magalhaes, I. S., Carleton, K. L., Mrosso, H. D. J., Miyagi, R., … Okada, N. (2008). Speciation through sensory drive in cichlid fish. Nature, 455(7213), 620–626. doi: 10.1038/nature07285

Sibeaux, A., Cole, G. L., & Endler, J. A. (2019). Success of the receptor noise model in predicting colour discrimination in guppies depends upon the colours tested. Vision Research, 159, 86–95. doi: 10.1016/j.visres.2019.04.002

Steiner, C. C., Römpler, H., Boettger, L. M., Schöneberg, T., & Hoekstra, H. E. (2009). The Genetic Basis of Phenotypic Convergence in Beach Mice: Similar Pigment Patterns but Different Genes. Molecular Biology and Evolution, 26(1), 35–45. doi: 10.1093/molbev/msn218

Stevens, M., Párraga, C. A., Cuthill, I. C., Partridge, J. C., & Troscianko, T. S. (2007). Using digital photography to study animal coloration. Biological Journal of the Linnean Society, 90(2), 211–237. doi: 10.1111/j.1095-8312.2007.00725.x

Stram, Daniel O., & Lee, J. W. (1994). Variance Components Testing in the Longitudinal Mixed Effects Model. Biometrics, 50(4), 1171–1177. JSTOR. doi: 10.2307/2533455

Stuart, Y. E., Veen, T., Weber, J. N., Hanson, D., Ravinet, M., Lohman, B. K., … Bolnick, D. I. (2017). Contrasting effects of environment and genetics generate a continuum of parallel evolution. Nature Ecology & Evolution, 1(6), 158. doi: 10.1038/s41559-017-0158

Suk, H. Y., & Neff, B. D. (2009). Microsatellite genetic differentiation among populations of the Trinidadian guppy. Heredity, 102(5), 425–434. doi: 10.1038/hdy.2009.7

Tao, L., DeRosa, A. M., White, T. W., & Valdimarsson, G. (2010). Zebrafish cx30.3: Identification and Characterization of a Gap Junction Gene Highly Expressed in the Skin. Developmental Dynamics?: An Official Publication of the American Association of Anatomists, 239(10), 2627–2636. doi: 10.1002/dvdy.22399

The Gene Ontology Consortium. (2019). The Gene Ontology Resource: 20 years and still GOing strong. Nucleic Acids Research, 47(D1), D330–D338. doi: 10.1093/nar/gky1055

Tripathi, N., Hoffmann, M., Willing, E.-M., Lanz, C., Weigel, D., & Dreyer, C. (2009). Genetic linkage map of the guppy, Poecilia reticulata, and quantitative trait loci analysis of male size and colour variation. Proceedings. Biological Sciences, 276(1665), 2195– 2208. doi: 10.1098/rspb.2008.1930

Troscianko, J., & Stevens, M. (2015). Image calibration and analysis toolbox – a free software suite for objectively measuring reflectance, colour and pattern. Methods in Ecology and Evolution, 6(11), 1320–1331. doi: 10.1111/2041-210X.12439

van den Berg, C. P., Troscianko, J., Endler, J. A., Marshall, N. J., & Cheney, K. L. (2020). Quantitative Colour Pattern Analysis (QCPA): A comprehensive framework for the analysis of colour patterns in nature. Methods in Ecology and Evolution, 11(2), 316– 332. doi: 10.1111/2041-210X.13328

Vorobyev, M., & Osorio, D. (1998). Receptor noise as a determinant of colour thresholds. Proceedings. Biological Sciences, 265(1394), 351–358. doi: 10.1098/rspb.1998.0302

Wang, F., Wolfson, S. N., Gharib, A., & Sagasti, A. (2012). LAR receptor tyrosine phosphatases and HSPGs guide peripheral sensory axons to the skin. Current Biology: CB, 22(5), 373–382. doi: 10.1016/j.cub.2012.01.040

Weadick, C. J., Loew, E. R., Rodd, F. H., & Chang, B. S. W. (2012). Visual pigment molecular evolution in the Trinidadian pike cichlid (Crenicichla frenata): a less colorful world for neotropical cichlids? Molecular Biology and Evolution, 29(10), 3045–3060. doi: 10.1093/molbev/mss115

Whitlock, M. C., & Guillaume, F. (2009). Testing for Spatially Divergent Selection: Comparing QST to FST. Genetics, 183(3), 1055–1063. doi: 10.1534/genetics.108.099812

Willing, E.-M., Bentzen, P., van Oosterhout, C., Hoffmann, M., Cable, J., Breden, F., … Dreyer, C. (2010). Genome-wide single nucleotide polymorphisms reveal population history and adaptive divergence in wild guppies. Molecular Ecology, 19(5), 968–984. doi: 10.1111/j.1365-294X.2010.04528.x

Xu, Y., & Fisher, G. J. (2012). Receptor type protein tyrosine phosphatases (RPTPs) – roles in signal transduction and human disease. Journal of Cell Communication and Signaling, 6(3), 125–138. doi: 10.1007/s12079-012-0171-5

Young, M. J., Simmons, L. W., & Evans, J. P. (2011). Predation is associated with variation in colour pattern, but not body shape or colour reflectance, in a rainbowfish (Melanotaenia australis). The Journal of Animal Ecology, 80(1), 183–191. doi: 10.1111/j.1365-2656.2010.01759.x

