## Appendix A for "Sensory-based quantification of male colour patterns in Trinidadian guppies reveals nonparallel phenotypic evolution across an ecological transition in multivariate trait space"

**Title:** Validation of phenotyping approach using captive-bred paternal sibships

**Rationale**

Before attempting to characterise putatively genetic differences in male phenotypes among-populations, we first tested whether the QCPA phenotyping method could reliably detect and characterise genetic variation using a sample of males from known paternal sibships. Male coloration and pattern traits are known to be at least partly Y-linked (Winge and Ditlevsen, 1947; Haskins et al., 1961). Thus, if the QCPA method provides a robust strategy for characterising (genetic) variation we predict high family-level repeatabilities (or intra-class correlations) for traits quantified. Note that while the presence or absence of body pattern elements is highly conserved with patrilineal lineages (figure A1), continuous traits are broadly expected to be subject to environmental and autosomal genetic effects too. Thus we do not expect a complete absence of within-family variation in QCPA derived trait.

**Methods**

Color pattern variation was opportunistically measured in fish from some paternal sibships produced for a different study (Bergero et al., 2019). In brief, after importing wild fish to the UK, small breeding group groups consisting of one male (sire) with distinct color patterns and several virgin females (dams 2-3) were isolated in an enriched (e.g. plants and substrate) 28L tank. Sires were haphazardly chosen from a few wild populations (QL, PM, GH, AL, QH, YH) representing both high and low predation regimes, and allowed to mate naturally with females. Females were from the same population in most, but not all cases. Offspring produced were separated from parental fish and raised to sexual maturity (as assessed from external morphology). Once male color patterns were fully expressed (2-3 months), males were photographed. In total the sample contained 137 individuals that mainly included male offspring from 13 families (Table A1).

*Photography of live animals*

Each guppy was individually placed in a custom designed UV-transparent (UV-transparent PMMA) and water filled chamber (50 L x 26 W x 30 mm H). The chamber contained an adjustable white PTFE background with a 1 cm scale bar and 95% and 5% reflectance standards were placed. We photograph unsedated live fish to maximize ecological relevance as far as possible (e.g. avoiding known expansion of melanophore cells that occurs with anaesthesia under MS222). Our setup also eliminates unnatural specular reflectance from the surface of the fish when removed from water.

Once each fish became still in the chamber, its left side was photographed. We captured the same image of the whole body of the fish in both visible and UV-ranges. Raw photographs were taken with a Samsung NX1000 camera converted to full spectrum, fitted with a Nikkor EL 80mm lens, capable of capturing colour patterns in both the human-visible and UV spectrum ranges (Troscianko and Stevens 2015). The lens was fitted with two sliding filters—a Baader UV-IR blocking filter (transmitting 420-680nm) and a Baader Venus-U UV pass filter (transmitting 320-380nm), allowing rapid capture of both images (within 1 sec). The chamber was illuminated from above by an Iwasaki eyeColour bulb simulating D65 lighting, with UVA emissions increased by removal of its UV-protective coating.

*QCPA Pipeline for phenotyping colour pattern variation*

RAW images of individual fish were analysed using the QCPA’s toolbox plug-ins (van den Berg *et al.* 2019) in ImageJ (Rasband 1997). UV and visible images were first combined and normalised using he 95% and 5% reflectance standards. All normalized images, in which the body (excluding the head, tail or fins) was selected as the region of interest (ROI), were mapped to the visual sensitivity the guppy (Kawamura *et al.* 2016; Kranz *et al.* 2018; Kranz *et al.* 2018) and the pike cichlid (Weadick *et al.* 2015) to obtain estimated photon catch values for global colour patch on the male body. The guppy retina possesses nine opsin genes, made up of four groups of photoreceptor cone types sensitive to UV (UVS), short (SWS), medium (MWS), and long (LWS) wavelength range (Kawamura *et al.* 2016). Following Kranz *et al.*(2018), spectral sensitivities (_max_) were selected based on microspectrophometric and opsin gene expression data, and include 359 nm (UVS), 408 nm (SWS), 465 nm (MWS) and 560 nm (LWS). The spectral sensitivity for the guppy double cones (or luminance) was based on the summation of the MWS and LWS spectra. On the other hand, the pike cichlid possesses three types of cones (SWS, MWS, LWS), with _max_ at 480 nm, 547 nm and 614 nm, respectively (Weadick *et al.* 2012).

Using QCPA, we quantified the body pattern conspicuousness using boundary strength analysis (BSA) (Endler *et al.* 2018) available in the Batch Multispectral Image Analysis option of the toolbox. In brief, the cone-catch image was first clustered using the “RNL clustering” function, breaking the image down into a discrete number of different colours based on the receiver’s colour vision and spatial acuity. The boundary strength analysis (BSA) was then used to describe the “saliency” of colour patterns. This analysis calculates the contrast between all pairs of colour patches in the image (e.g. chromaticity, colour (S) or luminance contrast), and creates a weighted average based on the degree to which each pair of patches are adjacent to each-other in the image. Mean values for both chromatic and achromatic aspects of body pattern were independently calculated, where high mean values typically indicate more stimulating and salient signals to the receiver (Endler *et al.* 2018). All colour perception metrics were based on plausible viewing distances from which a conspecific guppy or predator would assess a mate or attack a prey, respectively (Endler 1991; pike cichlid: 110mm, R. Heathcote, personal communication), the cone ratio abundance of each perceiver (guppy: 1:1:2:2 for UVS, SWS, MWS, LWS; Long, 1993; Laver and Taylor 2011; cichlid: 1:2:2 for SWS, MWS, LWS; Kröger *et al.* 1999), and visual acuity (guppy: 1.5 cycles per degree (cpd); Long 1993; cichlid: 12.6 cpd, based on Caves *et al.* 2016). We extracted and used for downstream analyses RNL-estimated means of saturation *sat_S_*, luminance boundary strength *lum_S_*, chromaticity boundary strength *chrom_S_* and chromaticity boundary strength variation *CoV_S_.*

*Statistical Analyses*

A univariate linear mixed model (LMM) was then fitted to each response variable with *family* identity as a random effect, and *cross* type as a fixed factor denoting whether dams were from the same population as the sire. No effect of *cross* was hypothesised or anticipated and it was included simply as statistical control for any variance generated. For each model, the among-family variance estimate (V_F_) was divided by the total variance conditional on fixed effects (calculated as the sum of V_F_ and within-family (residual) variance V_R_) to estimate the family level repeatability (R_F_). and likelihood ratio tests (LRT) were used to test the significance of the random effect variance by comparing full and reduced models. We assume that twice the difference in model log-likelihoods is distributed as a 50:50 mix of χ^2^_0_ and χ^2^_1_. Additionally, for each pair of homologous response variables derived using the conspecific and predator vision models (e.g., *chrom_ΔS_* under the conspecific and predator vision models) we also fitted a bivariate mixed model to estimate the family-level correlation (r_F_). This model was compared (by LRT on 1DF) to a constrained model in we added a constraint of r_F_=+1. This was to provide a simple evaluation of whether traits defined under the two vision models are likely to provide useful and distinct information about genetic differentiation (R_F_>0 for both traits and r_F_<+1). Mixed models were fit by REML in ASReml-R in R.

**Results**

Family-level descriptive statistics (means and standard deviations) on the QCPA-derived traits are shown in Table A2. Under the guppy vision model trait means were typically high (Mean_chrom_ = 2.39 ± 0.54, Mean_lum_ = 2.19 ± 0.69 e.g., greater than 1.5 for both chromatic and achromatic metric) and comparable to threshold values in Endler et al. (2018). Conpicuousness based on chromaticity was generally lower (Mean_chrom_ = 1.67 ± 1.26, Mean_lum_ = 2.86 ± 1.11), though not for luminance, under the cichlid vision model.

Plotting trait pairs within each vision model revealed evidence of phenotyping clustering by family (Figure A2). More formally, univariate mixed models confirmed significant among-family variance for all eight response variables. Estimated among-family repeatabilities (R_F_) ranging from 35% (chrom_ΔS_ under the guppy vision model) to 84% (chrom_ΔS_ under the cichlid vision model) with a median of 48.5%; Table A3). Family level correlations between homologous traits defined under different vision models ranged from 0.08 to 0.80 and were all significantly less than +1, except for the coefficient of covariation (Table A3).

**Conclusions**

The significant, moderate to high estimates of R_F_ confirm that the phenotyping approach used is able to detect among family differences that are known to arise from genetic (and at least partially Y-linked) factors. Furthermore, we find that while family effects are significant on all traits extracted under both vision models, the estimated cross-model family correlations (r_F_) for homologous traits are significantly less than 1, except for the coefficient of variation. Thus traits can generally be viewed as genetically variable aspects of body colour and pattern, but also highlights that homologous traits defined under the guppy and cichlid vision models are themselves genetically distinct. It does not necessarily follow that patterns of genetic covariation among families will be recapitulated among populations. Nor is it true that all among-population phenotypic variance must have a genetic basis (as plasticity will certainly contribute to genetically complex traits). However, these results do demonstrate the validity of the phenotyping strategy for genetic investigation of the among-population divergence, and highlight the value of considering both vision system models.

**References**

Bergero R, Gardner J, Bader B, Yong L, Charlesworth D. (2019). Exaggerated heterochiasmy in a fish with sex-linked male coloration polymorphisms. *Proceedings of the National Academy of Sciences USA* 116(14) 6924-6931.

Caves, E. M., Sutton, T. T., & Johnsen, S. (2017). Visual acuity in ray-finned fishes correlates with eye size and habitat. *The Journal of Experimental Biology*, *220*(Pt 9), 1586–1596. doi: 10.1242/jeb.151183

Endler, J. A. (1991). Variation in the appearance of guppy color patterns to guppies and their predators under different visual conditions. *Vision Research*, *31*(3), 587–608. doi: 10.1016/0042-6989(91)90109-i

Endler, J. A., Cole, G. L., & Kranz, A. M. (2018). Boundary strength analysis: Combining colour pattern geometry and coloured patch visual properties for use in predicting behaviour and fitness. *Methods in Ecology and Evolution*, *9*(12), 2334–2348. doi: 10.1111/2041-210X.13073

Kawamura, S., Kasagi, S., Kasai, D., Tezuka, A., Shoji, A., Takahashi, A., Imai, H., Kawata, M. (2016). Spectral sensitivity of guppy visual pigments reconstituted in vitro to resolve association of opsins with cone cell types. *Vision Research*, *127*, 67–73. doi: 10.1016/j.visres.2016.06.013

Kranz, A. M., Cole, G. L., Singh, P., & Endler, J. A. (2018). Colour pattern component phenotypic divergence can be predicted by the light environment. *Journal of Evolutionary Biology*, *31*(10), 1459–1476. doi: 10.1111/jeb.13342

Kranz, A. M., Forgan, L. G., Cole, G. L., & Endler, J. A. (2018). Light environment change induces differential expression of guppy opsins in a multi-generational evolution experiment. *Evolution* 72(8), 1656-1676. doi: 10.1111/evo.13519

Kröger, R. H. H., Bowmaker, J. K., Wagner, H. J. (1999). Morphological changes in the retina *Aequidens pulcher* (Cichlidae) after rearing in monochromatic light. *Vision Research*  2441-2448.

Laver, C. R. J., & Taylor, J. S. (2011). RT-qPCR reveals opsin gene upregulation associated with age and sex in guppies (Poecilia reticulata) - a species with color-based sexual selection and 11 visual-opsin genes. *BMC Evolutionary Biology*, *11*, 81. doi: 10.1186/1471-2148-11-81

Rasband, W. S. (1997). *ImageJ*. Bethesda, Maryland, USA: U. S. National Institutes of Health. Retrieved from <https://imagej.nih.gov/ij/>

Troscianko, J., & Stevens, M. (2015). Image calibration and analysis toolbox – a free software suite for objectively measuring reflectance, colour and pattern. *Methods in Ecology and Evolution*, *6*(11), 1320–1331. doi: 10.1111/2041-210X.12439

Weadick, C. J., Loew, E. R., Rodd, F. H., & Chang, B. S. W. (2012). Visual pigment molecular evolution in the Trinidadian pike cichlid (*Crenicichla frenata*): a less colorful world for neotropical cichlids? *Molecular Biology and Evolution*, *29*(10), 3045–3060. doi: 10.1093/molbev/mss115

| **Table A1**. Sample of lab-raised families and guppies used for phenotyping validation | | | |
| --- | --- | --- | --- |
| family name | N | Site Origin | Predation regime |
| AHP:1 | 10 | AH | high |
| ALP:1 | 12 | AL | low |
| ALP:2 | 19 | AL | low |
| ALP:B1 | 5 | AL | low |
| GHP:1 | 7 | GH | high |
| GHP:2 | 10 | GH | high |
| GLP:B1 | 12 | GL1 | low |
| PML:1 | 12 | PML | low |
| PML:2 | 11 | PML | low |
| QHP:1 | 14 | QH | high |
| QLP:1 | 8 | QL | low |
| QLP:2 | 12 | QL | low |
| YHP:1 | 5 | YH | high |

| **Table A2**. Summary statistics mean and SD (in parentheses) by family under conspecific (guppy) and predator (cichlid) vision models | | | | | | | | |
| --- | --- | --- | --- | --- | --- | --- | --- | --- |
| family | Guppy vision model | | | | Cichlid vision model | | | |
|  | *chrom_DS_* | *CoV_DS_* | *lum_DS_* | *sat_DS_* | *chrom_DS_* | *CoV_DS_* | *lum_DS_* | *sat_DS_* |
| AHP:1 | 3.42 (0.87) | 0.58 (0.11) | 0.34 (0.04) | 1.06 (0.36) | 5.5 (1.21) | 0.52 (0.25) | 0.62 (0.13) | 1.55 (0.17) |
| ALP:1 | 1.5 (0.47) | 0.78 (0.09) | 0.1 (0.02) | 1.66 (0.28) | 0.83 (0.37) | 1.12 (0.25) | 0.26 (0.10) | 2.58 (0.98) |
| ALP:2 | 2.91 (1.00) | 0.88 (0.32) | 0.26 (0.09) | 2.12 (0.41) | 1.69 (0.58) | 1.12 (0.26) | 0.37 (0.11) | 4.27 (1.62) |
| ALP:B1 | 1.79 (0.30) | 0.78 (0.13) | 0.14 (0.06) | 1.88 (0.17) | 0.63 (0.30) | 1.17 (0.24) | 0.19 (0.08) | 3.25 (0.35) |
| GHP:1 | 2.46 (0.30) | 0.68 (0.11) | 0.32 (0.05) | 3.4 (0.65) | 1.36 (0.34) | 1 (0.15) | 0.35 (0.04) | 1.62 (0.42) |
| GHP:2 | 2.03 (0.31) | 0.67 (0.12) | 0.2 (0.06) | 2.81 (0.40) | 0.55 (0.14) | 0.75 (0.50) | 0.18 (0.05) | 2.73 (0.77) |
| GHP:B1 | 1.82 (0.54) | 0.86 (0.17) | 0.15 (0.05) | 2.06 (0.42) | 1.86 (1.03) | 1.2 (0.46) | 0.36 (0.14) | 2.2 (0.68) |
| PML:1 | 2.24 (0.37) | 1.1 (0.13) | 0.2 (0.04) | 2.18 (0.37) | 1.53 (0.22) | 1.16 (0.09) | 0.34 (0.04) | 4.1 (0.72) |
| PML:2 | 2.67 (0.48) | 0.95 (0.21) | 0.24 (0.05) | 2.01 (0.36) | 1.59 (0.33) | 1.09 (0.19) | 0.35 (0.06) | 3.92 (0.65) |
| QHP:1 | 2.24 (0.48) | 0.42 (0.14) | 0.17 (0.04) | 1.19 (0.14) | 2.13 (0.34) | 0.51 (0.11) | 0.38 (0.03) | 0.71 (0.29) |
| QLP:1 | 2.45 (0.32) | 0.7 (0.14) | 0.28 (0.09) | 2.53 (0.62) | 1.83 (0.54) | 0.97 (0.33) | 0.38 (0.09) | 3.53 (0.91) |
| QLP:2 | 2.59 (0.36) | 0.67 (0.07) | 0.34 (0.05) | 3.18 (0.71) | 1.59 (0.33) | 1.08 (0.20) | 0.35 (0.05) | 3.9 (0.83) |
| YHP:1 | 3 (0.52) | 0.58 (0.13) | 0.18 (0.03) | 2.42 (0.53) | 0.7 (0.17) | 0.99 (0.20) | 0.21 (0.03) | 2.92 (0.38) |

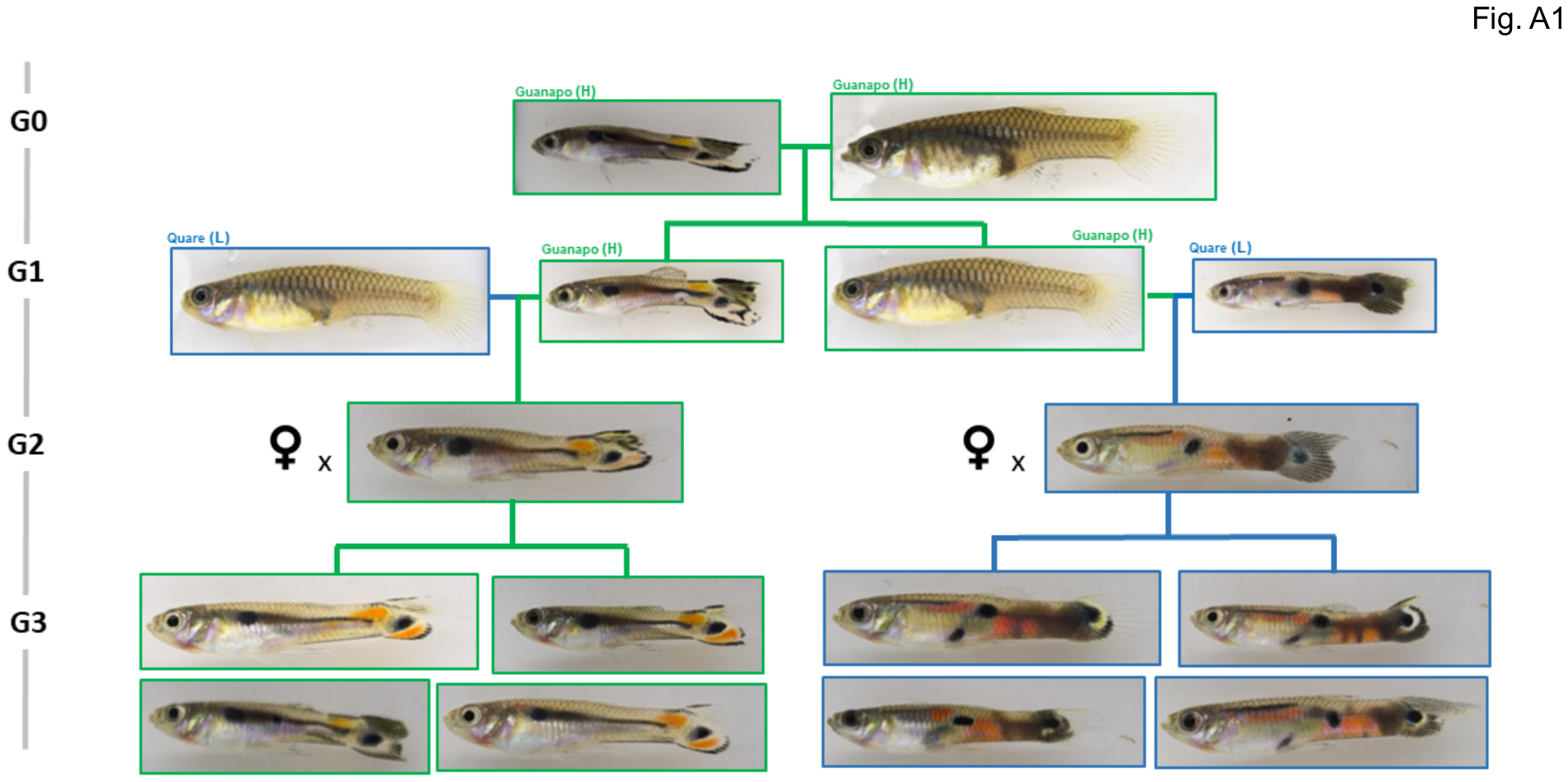

**Figure A1.** An example four generation pedigree in which conservation of coloration and pattern is visibly conserved down patrilines from grandsires to male grand-offspring.

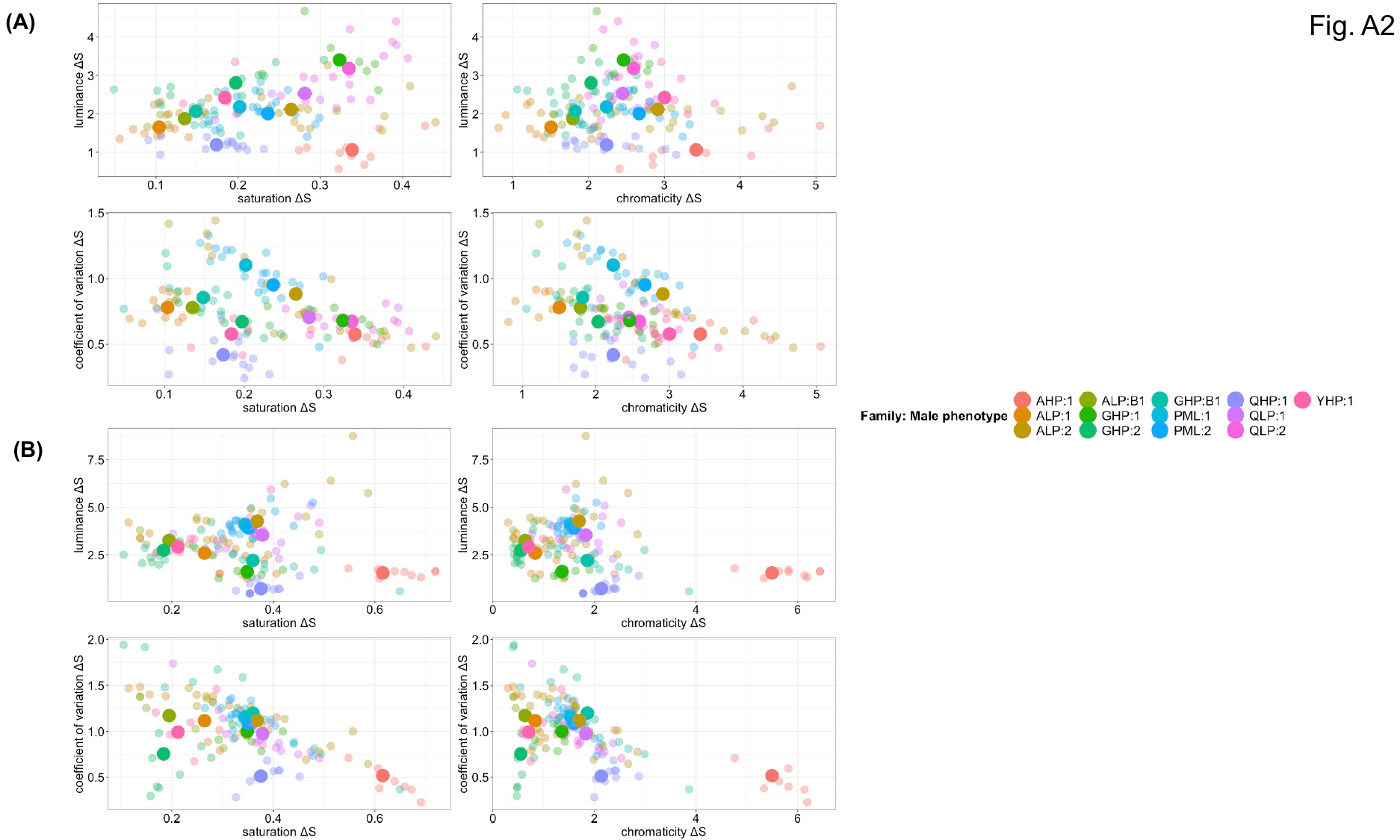

**Figure A2.** Pairwise plots of colour traits showing individual (small point) and family mean (large point with bars depicting SE) phenotypes for 13 lab-bred families. (A) depicts traits measured using the guppy vision model, while (B) are using the cichlid vision model. The plots represent the relationship of saturation and chromaticity with luminance and the coefficient of variation.
